# FALCON systematically interrogates free fatty acid biology and identifies a novel mediator of lipotoxicity

**DOI:** 10.1101/2023.02.19.529127

**Authors:** Nicolas Wieder, Juliana Coraor Fried, Choah Kim, Eriene-Heidi Sidhom, Matthew R. Brown, Jamie L. Marshall, Carlos Arevalo, Moran Dvela-Levitt, Maria Kost-Alimova, Jonas Sieber, Katlyn R. Gabriel, Julian Pacheco, Clary Clish, Hamdah Shafqat Abbasi, Shantanu Singh, Justine Rutter, Martine Therrien, Haejin Yoon, Zon Weng Lai, Aaron Baublis, Renuka Subramanian, Ranjan Devkota, Jonnell Small, Vedagopuram Sreekanth, Myeonghoon Han, Donghyun Lim, Anne E. Carpenter, Jason Flannick, Hilary Finucane, Marcia C. Haigis, Melina Claussnitzer, Eric Sheu, Beth Stevens, Bridget K. Wagner, Amit Choudhary, Jillian L. Shaw, Juan Lorenzo Pablo, Anna Greka

## Abstract

Cellular exposure to free fatty acids (FFA) is implicated in the pathogenesis of obesity-associated diseases. However, studies to date have assumed that a few select FFAs are representative of broad structural categories, and there are no scalable approaches to comprehensively assess the biological processes induced by exposure to diverse FFAs circulating in human plasma. Furthermore, assessing how these FFA-mediated processes interact with genetic risk for disease remains elusive. Here we report the design and implementation of FALCON (Fatty Acid Library for Comprehensive ONtologies) as an unbiased, scalable and multimodal interrogation of 61 structurally diverse FFAs. We identified a subset of lipotoxic monounsaturated fatty acids (MUFAs) with a distinct lipidomic profile associated with decreased membrane fluidity. Furthermore, we developed a new approach to prioritize genes that reflect the combined effects of exposure to harmful FFAs and genetic risk for type 2 diabetes (T2D). Importantly, we found that c-MAF inducing protein (CMIP) protects cells from exposure to FFAs by modulating Akt signaling and we validated the role of CMIP in human pancreatic beta cells. In sum, FALCON empowers the study of fundamental FFA biology and offers an integrative approach to identify much needed targets for diverse diseases associated with disordered FFA metabolism.

**Highlights:**

- FALCON (Fatty Acid Library for Comprehensive ONtologies) enables multimodal profiling of 61 free fatty acids (FFAs) to reveal 5 FFA clusters with distinct biological effects
- FALCON is applicable to many and diverse cell types
- A subset of monounsaturated FAs (MUFAs) equally or more toxic than canonical lipotoxic saturated FAs (SFAs) leads to decreased membrane fluidity
- New approach prioritizes genes that represent the combined effects of environmental (FFA) exposure and genetic risk for disease
- C-Maf inducing protein (CMIP) is identified as a suppressor of FFA-induced lipotoxicity via Akt-mediated signaling

## Introduction

Obesity promotes and exacerbates a growing burden of chronic diseases including type 2 diabetes (T2D), cancer, cardiovascular disease and neurodegeneration (Pillon et al. 2021; Rathmell 2021; Yoon et al. 2021). Obesity interacts with underlying genetic risk to drive this unprecedented health crisis; however, at present we do not understand the molecular mechanisms underpinning the complex interplay between genetic and environmental factors (Pillon et al. 2021). Genome wide association studies (GWAS) are powerful in identifying genetic risk factors but are limited by challenges in prioritizing disease-relevant loci (Pillon et al. 2021; Uffelmann et al. 2021). Metabolomic and lipidomic studies have linked aberrant levels of circulating lipids to metabolic diseases (Musunuru and Kathiresan 2019, 2016). However, the contribution of individual lipid species (i.e. triacylglycerols, lipoprotein particles, phospholipids, diacylglycerols and fatty acids) remains an area of active investigation (Farese et al. 2012; Jha et al. 2018; Imamura et al. 2017). Cellular models of fatty acid overload have been used to define “lipotoxicity” as the harmful effects of prolonged and increased exposure to specific lipids (Lytrivi, Castell, et al. 2020; Sharma and Alonso 2014), but the link between cellular exposure to diverse FFA species circulating in human blood and toxicity phenotypes remains poorly understood.

T2D, one of the most prevalent obesity-associated diseases, is characterized by insulin resistance in peripheral tissues and progressive pancreatic beta cell failure. Although liver, adipose and muscle tissue play a central role in insulin resistance (da Silva Rosa et al. 2020), T2D GWAS to date have highlighted genes critical to beta cell function (Petrie, Pearson, and Sutherland 2011; Pasquali et al. 2014; Udler et al. 2018; Parker et al. 2013). *In vitro*, beta cells are strongly affected by lipid overload; excess chronic fatty acid exposure disrupts their metabolic homeostasis and their capacity to secrete insulin, pushing them towards ER stress and cell death (Prentki and Madiraju 2012; Randle et al. 1963). Therefore, FFA studies in beta cells are centrally important to our understanding of the combined effects of excess FFA exposure and genetic risk for T2D.

Not all fatty acids produce the same cellular effects, and distinctions are often made based on their saturation level. Saturated fatty acids (SFAs) contain no double bonds in their carbon chains. In contrast, monounsaturated fatty acids (MUFAs) contain one double bond and polyunsaturated fatty acids (PUFAs) contain more than one double bond. At the cellular level, studies to date have primarily relied on the effects of a single SFA, palmitic acid (PA) (Palomer et al. 2018; Piccolis et al. 2019) to explore the role of FFAs in lipotoxicity. However, large-scale epidemiological studies, including the Framingham Heart Study, have shown an association between the abundance of TAG species composed of a wide spectrum of structurally diverse FFAs and the severity of metabolic diseases (Rhee et al. 2011), hinting at the importance of largely unexplored and clinically relevant FFA biology. Considering that (i) TAGs are hydrolyzed by lipases into FFAs (and glycerol) before uptake into peripheral cells (Saponaro et al. 2015), and (ii) there is a strong association between TAG composition and obesity-associated diseases such as T2D (Al-Sulaiti et al. 2018; Rhee et al. 2011), there is an urgent need to thoroughly interrogate the biological effects of FFAs across their structural spectrum.

Here we reveal FALCON (Fatty Acid Library for Comprehensive ONtologies), a cell-based platform for the unbiased, multimodal investigation of structurally diverse FFAs found in human plasma (Abdelmagid et al. 2015; Aubourg et al. 1993; Lust et al. 2021) by integrating transcriptomics, cell morphological features, lipidomics, FFA structural characterization, and functional profiling. Out of 61 FFAs, FALCON identified 20 FFAs that were toxic to beta cells, grouped into a cluster marked by a distinct transcriptomic signature. These lipotoxic FFAs could not be classified based on traditional structural designations such as degree of saturation. More than half (12/20) were MUFAs whose toxicity is not fully understood on a mechanistic level. Exposure to these MUFAs was associated with a distinct lipidomic profile and decreased membrane fluidity. Aiming to understand the combined effects of environmental exposure and genetic risk for T2D, we prioritized 25 genes that emerged from the overlap between genes differentially regulated after exposure to lipotoxic FFAs and genes nominated in a large-scale T2D GWAS. We thus identified *CMIP*, a gene with no prior mechanistic links to metabolic disease, and showed that it protects mouse and human pancreatic beta cells from excess fatty acid exposure. In sum, FALCON empowered a comprehensive, multiplexed query of FFA biology and revealed *CMIP* as a previously unrecognized suppressor of lipotoxicity.

## Results

### A systematic approach to study FFA biology

To enable the systematic study of FFAs, we needed to (i) curate a comprehensive set of FFAs beyond those traditionally used in targeted experiments (mostly PA and oleic acid (OA)), (ii) engineer an approach that would solve the difficulty of working with compounds of varying levels of hydrophobicity, and (iii) assess the biological effects of these FFAs in an unbiased fashion. FALCON fulfilled all three criteria, and we used it to systematically interrogate 61 structurally diverse FFAs (Table S1) using multiple high-content assays in a manner widely applicable to any cell type of choice. Most of the analyzed FFAs were readily detectable in both human and mouse blood (Table S1).

We performed proof of concept studies in MIN6 cells, a widely used mouse insulinoma pancreatic beta cell line that displays robust glucose-sensing and insulin secretion, and shares well-conserved molecular programs with human beta cells (Fang et al. 2019; Skelin, Rupnik, and Cencic 2010). We generated several datasets to deeply phenotype the effects of treating cells with each of the 61 FFAs including measures of cell viability/function, lipidomics, transcriptomics and unbiased cell morphology measurements (Fig. 1A, tier 1). We analyzed and integrated these diverse datasets using (i) Gene Set Enrichment Analysis (GSEA (Subramanian et al. 2005)) to identify alterations in cellular pathway activity; (ii) Molecular Operating Environment (MOE) software (Vilar, Cozza, and Moro 2008) to correlate FFA chemical structures with biological features, and (iii) Multimarker Analysis of GenoMic Annotation (MAGMA (de Leeuw et al. 2015)) ranking to incorporate genetic predisposition to human metabolic disease (Fig. 1A, Tier 2). Importantly, we validated our results from MIN6 cells in two human beta cell systems: iPSC derived pancreatic beta cells and acutely isolated human pancreatic islets from donors. Collectively, this multimodal approach allowed us to (i) group FFAs in an unbiased manner (i.e. remaining agnostic to structural features) solely based on similarity of biological readouts, (ii) identify previously unrecognized lipotoxic FFAs, and (iii) define toxicity phenotypes representative of these lipotoxic FFAs.

**Fig. 1.**
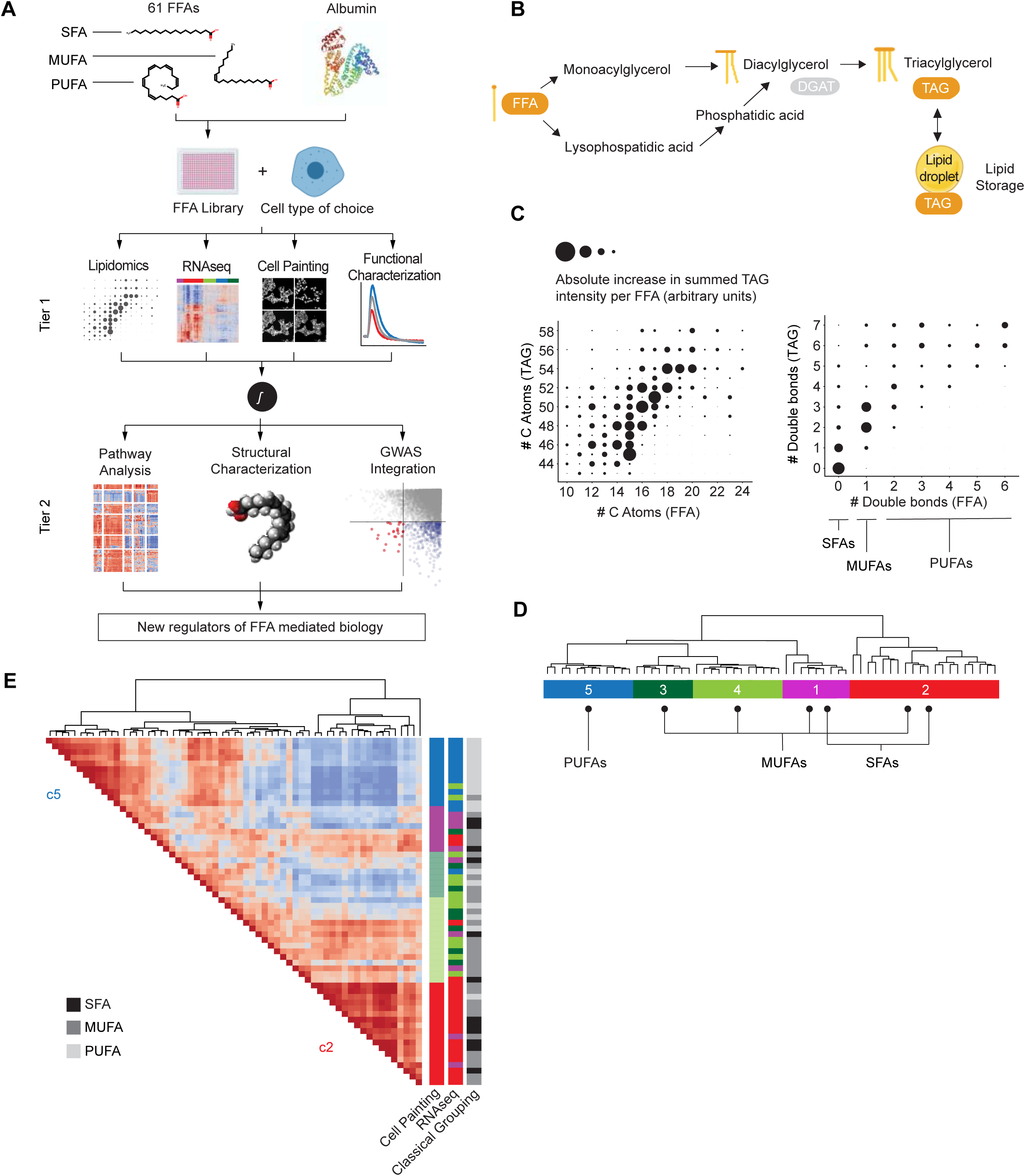
FALCON, a multiplexed platform for the systematic interrogation of structurally diverse FFAs defines 5 FFA clusters. (A) Analysis workflow for FALCON. SFA, saturated fatty acid; MUFA, monounsaturated fatty acid; PUFA, polyunsaturated fatty acid. In Tier 1, we use several methods, as listed, to characterize the cellular effects of each of 61 FFAs. In Tier 2, the Tier 1 datasets are integrated to reveal mechanistic insights. (B) Schematic of triacylglyceride (TAG) synthesis from FFAs. Shown here are the monoacylglycerol (top) and the glycerol-3-phosphate pathway (bottom). For simplicity, the acylation of dihydroxyacetone phosphate is not shown. (C) Qualitative correlation of structural features (number of C atoms, number of double bonds) of externally applied FFAs from the library (x-axis) versus structural features of endogenous TAGs (y-axis) measured by lipidomics. Distinct TAG profiles detected in cells treated with SFAs, MUFAs or PUFAs. (D) Five FFA clusters (c1-c5) were identified after hierarchical clustering of transcriptomic profiles derived from exposure to each of 61 FFAs (methods). (E) Cell Painting analysis of immunofluorescence images from cells exposed to each of 61 FFAs (methods) independently clustered together the FFAs transcriptomically assigned to c2 and separately the FFAs assigned to c5.

The crucial gating step in establishing FALCON was the ability to reproducibly and reliably deliver solvent-free BSA-bound FFAs to cells in multiwell plates at scale, using concentrations of magnitude similar to human blood (Abdelmagid et al. 2015; Aubourg et al. 1993; Tsoukalas et al. 2019; Mir et al. 2015; Mogilenko et al. 2019; Chen et al. 2020; Ubellacker et al. 2020; Lytrivi, Ghaddar, et al. 2020)). To achieve this goal, FFAs were dissolved in DMSO and non-covalently bound to FFA-free BSA at a 7:1 ratio (there are 7 FFA binding sites in human albumin (Petitpas et al. 2001; Curry et al. 1998; Bhattacharya, Grüne, and Curry 2000; Fujiwara and Amisaki 2008)). A centrifugal evaporation/concentration step removed water and DMSO to generate solvent-free crystals of BSA-bound FFAs (Fig. S1A). The crystals were dissolved in culture medium, which was then applied to cells (Fig. S1A). We confirmed FFA binding to BSA by differential scanning calorimetry (Michnik 2003)(Fig. S1B).

To assess the effects of FFA treatment on cellular lipid composition, we performed mass spectrometry-based lipidomics of cells exposed to each of the 61 FFAs (Fig. S1C). TAGs, which are synthesized from FFAs available within a cell (Coleman and Mashek 2011) (Fig. 1B), constituted the majority of detected lipid species and thus became the focus of our initial lipidomic analysis. We concluded that FFAs were likely incorporated into cellular TAGs, as evidenced by the correlation between FFA and TAG structural features, such as chain length and number of double bonds (Fig. 1C, Fig. S1D). In summary, FALCON assessed the biological effects of structurally diverse FFAs in a reliable, high-throughput, and quantitative manner.

### Identification of non-canonical FFA clusters

To comprehensively define FFA-induced cell states without prior assumptions about biological effects or FFA structural features, we generated transcriptomic profiles of MIN6 cells in response to each of 61 FFAs in the library (Fig. S1E). We detected induction of carnitine palmitoyltransferase 1 expression (CPT1A), the rate-limiting enzyme of FFA beta-oxidation (Thumelin et al. 1997), across the entire library, demonstrating successful FFA delivery and intracellular metabolism (Fig. S1F).

Hierarchical clustering of FFA-induced transcriptomes revealed 5 distinct “clusters” (c1- c5)(Fig. 1D, Fig. S2A,B). At the extremes, FFAs segregated by saturation and chain length: SFAs were major constituents of cluster 1 (c1) and also well represented in cluster 2 (c2; 8/20 c2 FFAs were SFAs), whereas cluster 5 (c5) was exclusively composed of PUFAs. In line with the traditional paradigm (Palomer et al. 2018), palmitic acid (lipotoxic FFA) and oleic acid (OA; lipotoxicity-protective FFA), clustered separately, in c2 and c3, respectively. Strikingly, MUFAs were distributed across four clusters c1 to c4 (Fig. 1D), indicating that the transcriptomic signatures classified some MUFAs as functionally similar to SFAs, and not to other MUFAs. These data argue against a simple grouping of FFAs based on saturation (SFA, MUFA, and PUFA) (Palomer et al. 2018) and suggest instead that traditional criteria do not adequately capture the observed transcriptomically-defined heterogeneity (Fig. S2C).

To probe cellular responses to FFA exposure using an independent method, we profiled the entire FFA library using Cell Painting, a high-content image analysis method that simultaneously measures several hundred cellular imaging features (Bray et al. 2016) (Fig. S2D). This morphologic analysis independently clustered together the SFAs and MUFAs transcriptomically assigned to c2 and separated them from the PUFAs in c5 (Fig. 1E). We thus concluded that the algorithms analyzing the cell imaging data were in agreement with the FFA clustering derived from transcriptomics with regard to the two most prominent clusters: c2 and c5 (Fig. 1D-E).

### Cellular transcriptomes define key biological responses to FFAs

To understand the basis behind the FFA clusters identified by FALCON, we applied Gene Set Enrichment Analysis (GSEA) (Subramanian et al. 2005) to each of the 61 FFA-induced transcriptomes using multiple gene set collections (MSigDB (Liberzon et al. 2015)). By looking at how multiple gene sets were enriched across the 5 clusters, we were able to categorize them into modules of gene expression (Fig. 2A; Methods). Crucially, this analysis ensured that our gene set annotation was comprehensive and not reliant on any single database of curated gene lists.

**Fig. 2.**
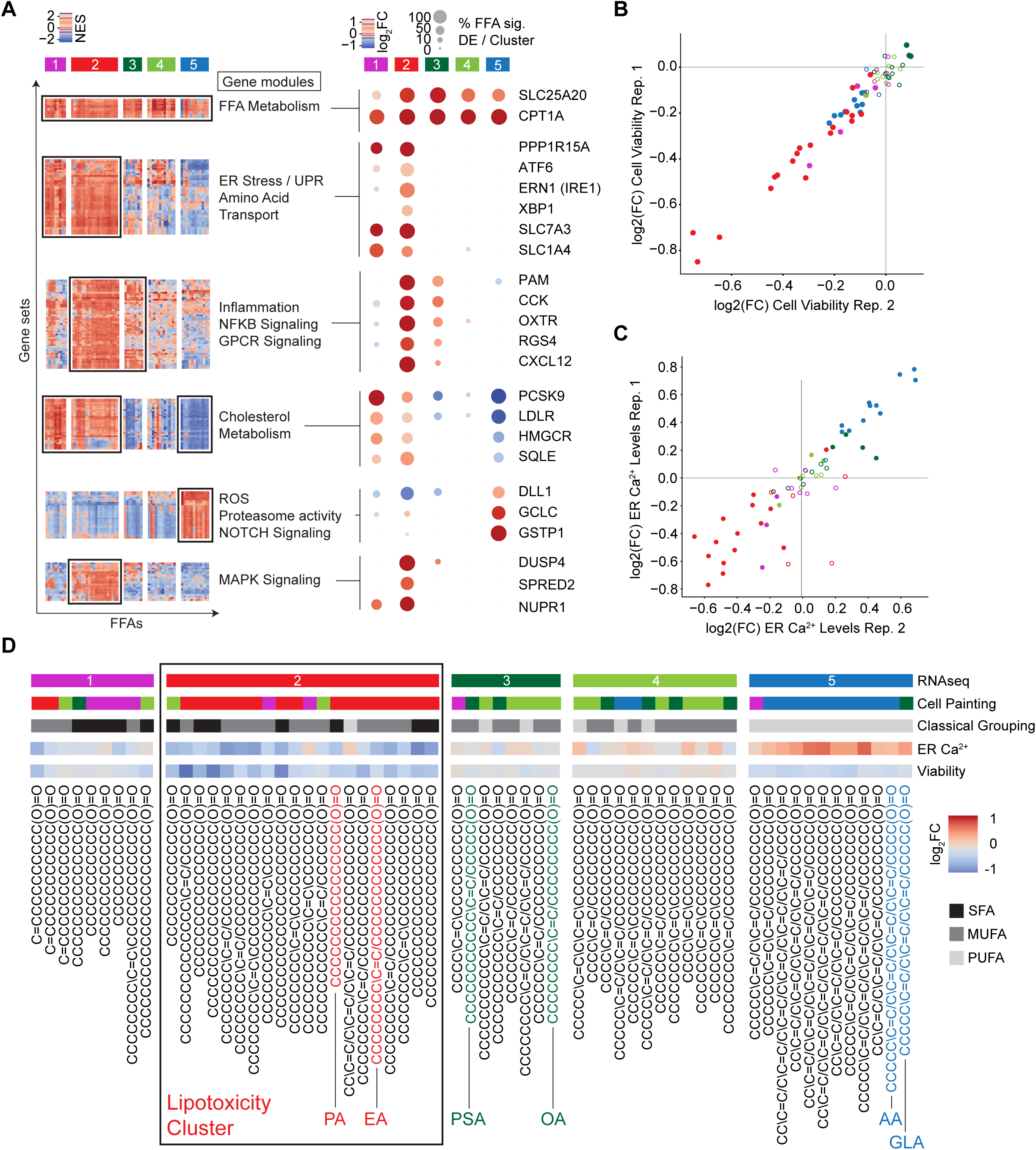
Transcriptomic analysis identifies key biological responses to FFAs, and functional assays validate novel FFA clustering. (A) Hierarchical clustering of a gene set enrichment matrix (based on normalized enrichment scores of gene sets, NES) revealed gene set modules of interest. Representative leading edge genes from each module are listed on the right. (B, C) Scatter plots of two independent replicates of cell viability (B) and ER Ca^2+^ level measurements (C)(n=5-7 replicates / FFA / screen). Closed dots represent FFAs that showed significant difference (p < 0.05, Bonferroni) from controls in both replicates, open dots represent non-significant FFAs in at least one replicate. Colors indicate corresponding FFA cluster membership (Fig. 1D). (D) Summary of functional assays. Top bar represents FFA clusters derived from transcriptomic analysis. Second color bar represents clusters derived from Cell Painting (cellular morphology) analysis (methods). Bar in grayscale indicates classical grouping of FFAs based on saturation level. The next two bars are heat maps displaying log2 fold changes of ER Ca^2+^ levels and cell viability, respectively. X-axis labels show FFA structure (in Simplified Molecular Input Line Entry System, SMILES). The box highlights the 20 FFAs in cluster 2 (c2) identified as the lipotoxicity cluster. Highlighted FFAs were chosen as cluster representatives for further downstream studies. PA, palmitic acid; EA, erucic acid; PSA, petroselenic acid; OA, oleic acid; AA, arachidonic acid; GLA: gamma- linoleic acid.

Since the identified gene modules served as the basis for the separation of diverse FFAs into five discrete clusters, we next sought to better understand the biological processes these modules represented. Consistent with exposure of cells to FFAs, a key module shared among all clusters represented pathways regulating FFA transport and metabolism, including upregulation of *SLC25A20* and *CPT1A* (core genes that encode the mitochondrial carnitine shuttle (Tonazzi et al. 2021; Longo, Frigeni, and Pasquali 2016). A second module, whose induction (positive normalized enrichment score (NES)) was detected specifically in c1 and c2, consisted of ER stress and unfolded protein response (UPR) genes (*PPP1R15A*, *ATF6*, *ERN1* (*IRE1*) and *XBP1;* (Schröder and Kaufman 2005)) (Fig. 2A). A third gene module comprised stress response pathways, NFkB signaling and inflammation (Fig. 2A) consistent with lipotoxicity (Hotamisligil 2010; Eguchi et al. 2012; Donath et al. 2013). Genes in the third module such as *PAM*, *CCK, OXTR*, *RGS4* and the chemokine, *CXCL12* were specifically upregulated in the c2 FFA cluster. Module four comprised pathways related to cholesterol metabolism (including key genes *PCSK9*, *LDLR*, *HMGCR* and *SQLE;* (Abifadel et al. 2003; Hobbs, Brown, and Goldstein 1992; Howe et al. 2017)), and module five consisted of programs related to proteasome activity and responses to reactive oxygen species (ROS) (upregulated genes *DLL1, GCLC, GSTP1* in the c5 cluster) (Fig. 2A). A sixth gene module was associated with MAPK signaling, a crucial regulator of cell proliferation and apoptosis that directly impacts insulin secretion in beta cells (Khoo et al. 2004). The MAPK signaling module was specifically upregulated in the c2 cluster (as reflected by the induction of genes such as *DUSP4*, *SPRED2* and *NUPR1* (Fig. 2A)). In sum, FALCON defined the biological processes that served as the basis for the separation of diverse FFAs into five previously unrecognized clusters. More specifically, the second, third and sixth modules pointed to the c2 cluster as a putative mediator of cell stress and lipotoxicity.

To explore the biological relevance and reproducibility of the newly defined c2 transcriptomic signature, we asked if differentially expressed genes from published lipotoxicity datasets in other systems might overlap with genes differentially expressed upon exposure to c2 FFAs. We found a distinct overlap of genes upregulated by c2 FFAs in MIN6 cells with (i) genes upregulated in the transcriptome of pancreatic beta cells isolated from mice fed a high fat diet (Dusaulcy et al. 2019) and (ii) genes upregulated in human islets exposed to PA (Lytrivi, Ghaddar, et al. 2020) (Fig. S2E,F; Methods). These analyses indicated that cellular responses to c2 FFAs were conserved and shared between mouse MIN6 cells and human beta cells *in vitro*, as well as mouse beta cells *in vivo*.

### Functional validation of FFA clustering

To functionally validate the transcriptome-derived clusters, we first measured cell viability in MIN6 cells exposed to each of the 61 FFAs (Fig. 2B). The c2 FFAs induced the most consistent and significant reduction in cell numbers among all clusters. Based on the observation that FFA-induced ER stress is associated with decreased ER Ca^2+^ levels and apoptosis (Marmugi et al. 2016; Fu et al. 2011; Orrenius, Zhivotovsky, and Nicotera 2003), we measured ER Ca^2+^ in a high-throughput fluorometric assay after exposure to each of the 61 FFAs (Fig. S3A, Methods). We detected decreased ER Ca^2+^ levels in cells treated with c2 FFAs and, to a smaller extent, c1 FFAs, thus functionally validating the ER stress signature derived from transcriptomics (Fig. 2C). In contrast, we detected a consistent increase in ER Ca^2+^ levels for cells exposed to c5 FFAs, while c3 and c4 FFAs did not alter ER Ca^2+^ levels (Fig. 2C). Collectively, these data showed that, in addition to SFAs and in contrast to c3 and c4 MUFAs, a specific subset of 12 MUFAs in the c2 cluster caused ER Ca^2+^ deficits and cell death (Fig. 2D). These results enhanced our interest into further understanding c2 FFAs, including c2 MUFAs, as mediators of lipotoxicity. More broadly, these data served as functional validation for FALCON in pancreatic beta cells.

### FALCON is applicable at scale to different cell types

Exposure to excess circulating FFAs has been implicated in diseases affecting many cells and organs in addition to pancreatic beta cells, including the kidney (Sun et al. 2020; Jang et al. 2020) and the brain (Yin 2022; Guttenplan et al. 2021; Ioannou et al. 2019). To assess the general versatility of FALCON, we tested two additional disease-relevant cell types: human kidney tubular epithelial cells, associated with kidney disease (Sun et al. 2020; Jang et al. 2020), and human iPSC-derived microglia, associated with neurodegenerative disease (Yin 2022; Tcw et al. 2022). In both cell types, c2 cluster FFAs were the most toxic, consistent with the results in beta cells (Fig. 3A-E). Of note, there were also some cell-type specific differences. In human kidney epithelial cells, c3-4 FFAs universally increased cell numbers, indicating that these FFAs may promote cell proliferation (Fig. 3A,D). In microglia, c5 PUFAs were more toxic than in beta cells and kidney epithelial cells, on par with c2 FFAs (Fig. 3A). High levels of PUFAs are associated with increased levels of ROS-induced damage due to lipid peroxidation leading to ferroptosis (W. S. Yang et al. 2016; Dixon et al. 2012; Ryan et al. 2023). Our data, the first comprehensive snapshot of the effect of diverse FFAs on human microglia, suggest that these critical neuro-immune cells may be particularly susceptible to PUFA-mediated injury (Fig. 3A), an intriguing finding with implications for neurodegenerative diseases (Bachiller et al. 2018). In sum, these studies demonstrated the utility of FALCON for multiple cell types of interest, and its potential to yield fundamental insights into FFA biology.

**Fig. 3.**
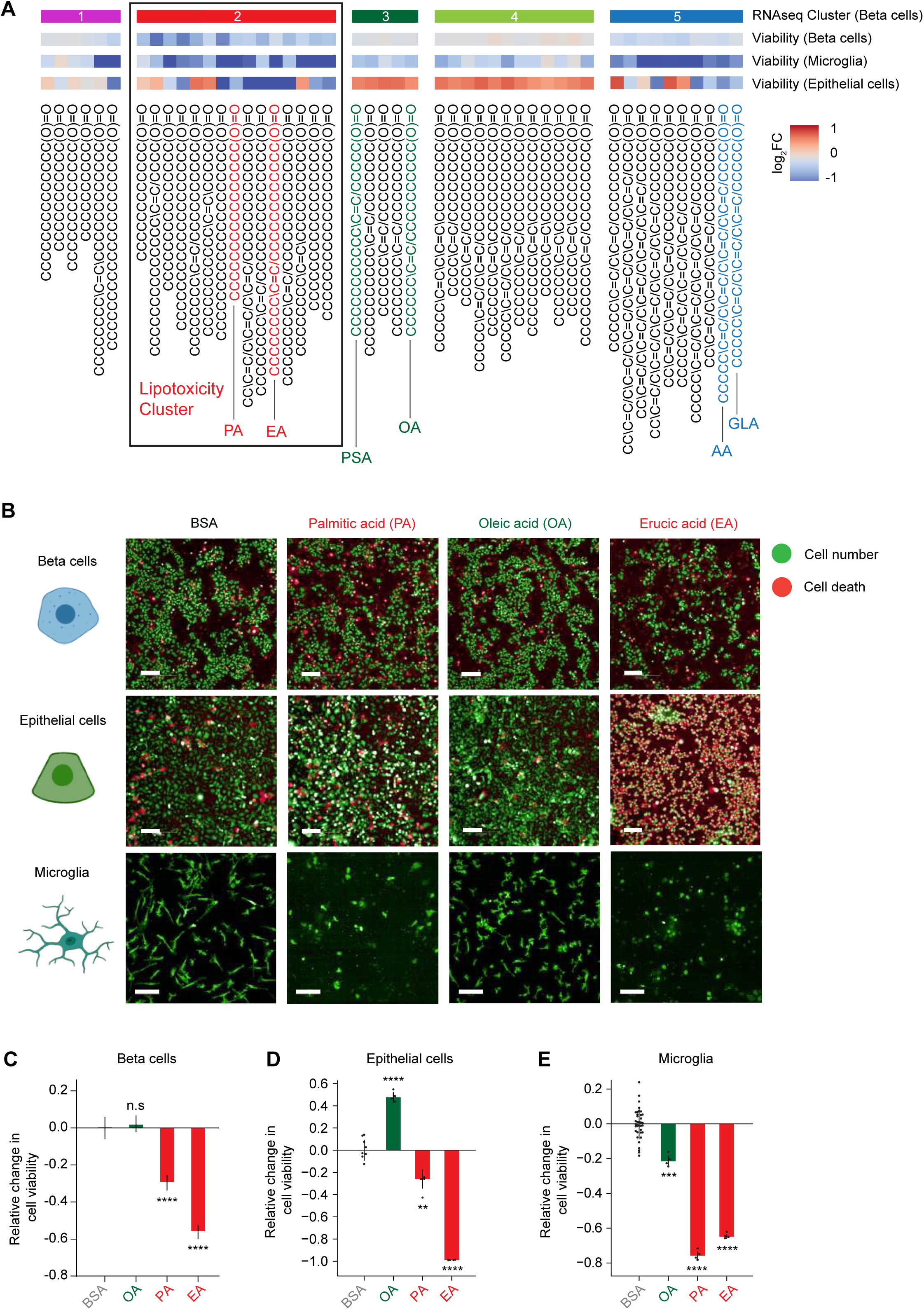
FALCON is applicable at scale to different cell types. (A) Heatmap showing comparison of viability changes across 3 different cell types: MIN6 pancreatic beta cells, human iPSC-derived microglia and human kidney tubular epithelial cells. c2 FFAs are toxic for all three cell types studied. (B) Representative images from all 3 cell types highlighting the toxicity of c2 FFAs including erucic acid (EA). EA induces cell death in beta cells (green, Hoescht), in microglia (green, GFP) and in kidney tubular epithelial cells (red, propidium iodide). Scale bars: 100 µm. (C-E) Bar plots indicate change in cell viability relative to BSA induced by OA, PA, or EA in each cell type after exposure to FFAs at 500 µM for 72 h (MIN6) (C), 500 µM for 15 h (epithelial cells) (D), or 250 µM for 24 h (microglia) (E). PA and EA consistently induce cell death across all three cell types as assessed by 1-way ANOVA followed by Dunnett’s test (\**p* < 0.05, \*\*\*\**p* < 0.0001). Data are mean ± SD.

### Cell biological hallmarks of lipotoxicity characterize the newly-defined c2 FFA cluster

To better understand the mechanisms underlying c2 FFA-induced toxicity, which was universally observed in beta cells, kidney epithelial cells and microglia, we took a closer look at the 8 SFAs and 12 MUFAs comprising this cluster (Fig. 2D, 3A) and focused on erucic acid (EA), a 22-carbon c2 MUFA. Although there is evidence that long chain MUFAs such as EA are toxic (Plötz et al. 2019), the mechanisms behind this toxicity are only beginning to be explored (Chen et al. 2020). In addition, previous *in vivo* studies had not clarified whether exposure to EA is harmful or not (Vogtmann et al. 1975; Aubourg et al. 1993). In our studies, despite its classification as a MUFA with a similar length to various c5 PUFAs, EA was distinctly clustered with lipotoxic c2 FFAs. To more deeply characterize EA, we examined it in comparison to (i) palmitic acid, a saturated c2 FFA, (ii) oleic acid (OA) and petroselinic acid (PSA), two MUFAs in the non-toxic c3 cluster, and (iii) arachidonic acid (AA) and gamma-linoleic acid (GLA), two long chain FFAs in the c5 PUFA cluster. Given the high disease burden from T2D and the availability of well-characterized cellular assays (as described for example in Figs. 1, 2), we focused our analysis on beta cells. Out of the six FFAs studied, only EA and PA consistently induced cell death (Fig. 4A).

**Fig. 4.**
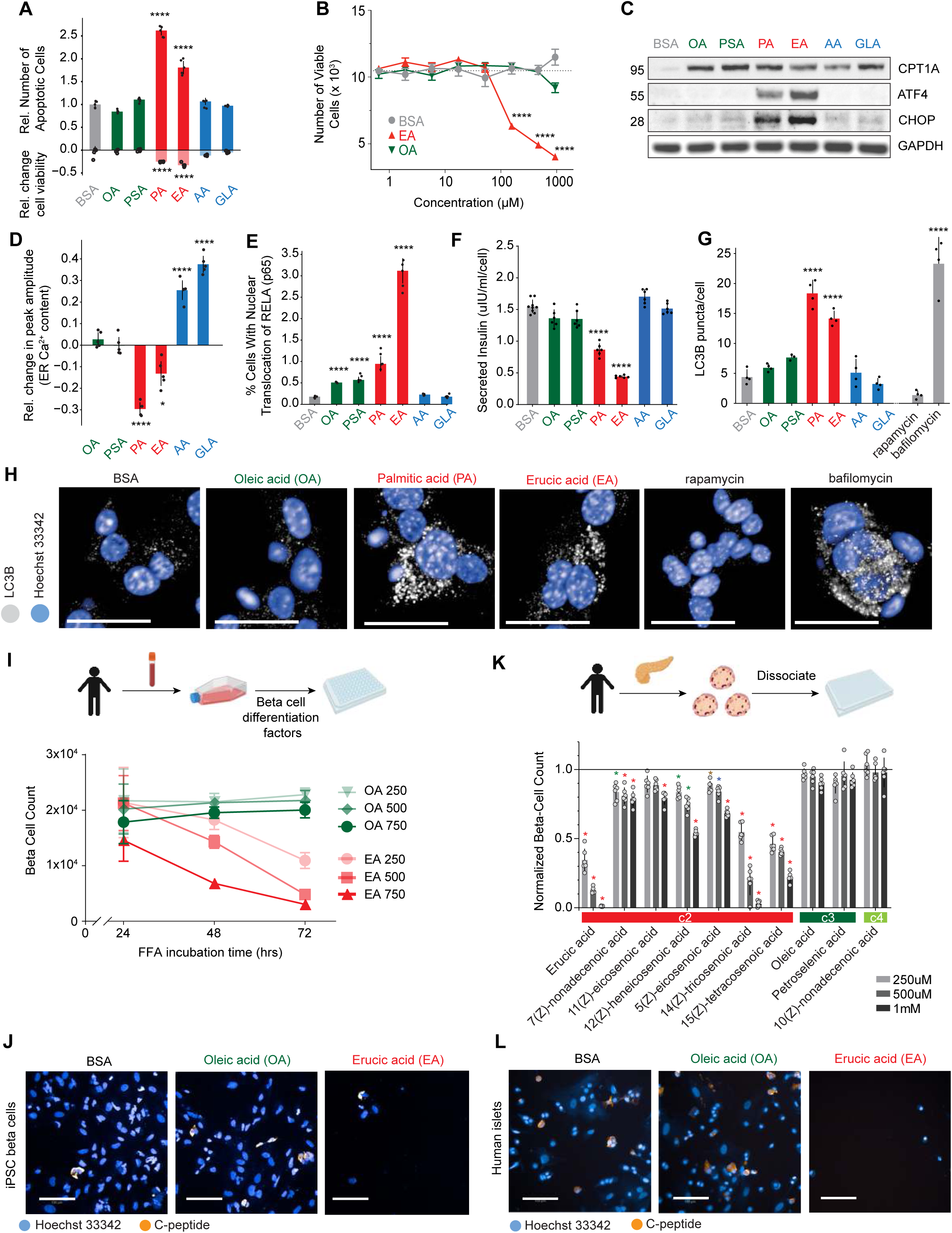
Cell biological hallmarks of lipotoxicity characterize the c2 FFA cluster. (A) Quantifica- tion of fluorescence imaging of cells treated with representative FFAs for 48 h. Apoptotic cells (positive y-axis) are measured by caspase activity. Dead cells are measured by propidium iodide positive nuclei (n = 5 wells). Reduction in cell viability (negative y-axis) is defined as the fraction of caspase positive and propidium iodide positive cells. Data are mean ± SD. Student’s t-test (two-sid- ed) ****p < 0.0001, corrected for multiple testing (Bonferroni). (B) Dose-response curve of live cell numbers after 65 h of EA or OA treatment compared to BSA. Live cell number is assessed by total number of Hoechst-positive nuclei that were negative for propidium iodide and for cleaved caspase 3/7. EA is toxic in a concentration dependent manner; OA is not toxic. (C) Western blots show ATF4 and CHOP induction by PA and EA (lipotoxic c2 FFAs). CPT1A induction is a control for intracellular FFA delivery. BSA, negative control. (D) Quantification of peak amplitude as readout for ER Ca2+ levels, relative to negative control (BSA). Data are mean ± SD. Student’s t-test (two-sided) *p < 0.05, ****p < 0.0001, corrected for multiple testing (Benjamini-Hochberg, entire FFA library). (E) Quantification of RELA translocation as percentage of total number of cells (t = 18 h, n = 5 replicates). Data are mean ± SD. Student’s t-test (two-sided) ****p < 0.0001, corrected for multiple testing (Bonferroni). (F) Glucose stimulated insulin secretion (GSIS) after FFA exposure ([FFA] = 500 µM, t = 24 h, n = 6 wells) was measured by ELISA. (G) Autophagosome formation was assessed by imaging LC3B puncta normalized to the number of total cells (t = 48 h, n = 4 wells). Data are mean ± SD. One-way ANOVA followed by Dunnett’s test. For A, D, E, G and H: bar color represents cluster identity (green, c3; red, c2; blue, c5; Fig. 1D). (H) Representative images of LC3B immunofluorescence (gray) in MIN6 cells treated with FFAs, BSA as negative control, or autophagy modulating drugs. Nuclei were detected by Hoechst (blue). c2 FFAs induce autophago- somes. Scale bars: 25 μm. (I) Number of iPSC-derived beta cells after exposure to EA or OA for 24, 48, or 72 h at 250 μM, 500 μM, or 750 μM. Only EA decreased cell count in a dose and time dependent manner. All EA conditions are significant per two-way ANOVA (p < 0.0001); all OA conditions are not significant. Data are mean ± SD, n = 4-6 wells/timepoint/FFA. (J) Representative images of iPSC-derived beta cells after treatment with BSA, EA, or OA for 48 h at 500 μM. Nuclei are marked by Hoechst and beta cell identity is marked by C-peptide staining (orange). Scale bars: 100 μm. (K) Quantification of C-peptide positive cells human beta cells dissociated from cadaveric primary islets normalized to the BSA control after exposure to each of 10 FFAs at 3 different con- centrations (t = 5 days, n = 6 wells). c2 cluster MUFAs decreased human beta cell viability in a dose-dependent manner. Data are mean ± SD. Multiple t-test with Bonferroni correction (gold: p<0.05, blue: p<0.01, green: p<0.001, red: p<0.0001). (L) Representative images of human islets after treatment with BSA, EA, or OA. Nuclei are marked by Hoechst, and beta cell identity is marked by C-peptide staining (orange). Scale bars: 100 μm. PA, palmitic acid; EA, erucic acid; PSA, petroselenic acid; OA, oleic acid; AA, arachidonic acid; GLA: gamma-linoleic acid.

Next we sought to explore the dose-dependent toxicity of EA, an FFA whose abundance in human plasma may be 10-200 times lower than PA (one of the most well-studied and abundant FFAs) (Abdelmagid et al. 2015; Aubourg et al. 1993; Lust et al. 2021). Recognizing that (1) plasma concentrations are not indicative of tissue concentrations (D. Zhang et al. 2019) and (2) FFA concentrations have not been measured in any tissue, including the pancreas, we tested a range of EA concentrations. We found that as low as ∼75-100 µM EA, a concentration previously measured in human plasma (Aubourg et al. 1993; Lust et al. 2021), induced high levels of cell death (Fig. 4B). These dose-response experiments confirmed that the biological effects we measured were physiologically relevant for PA and toxic MUFAs such as EA.

Sustained activation of the UPR has been linked to the toxicity induced by prolonged exposure to SFAs. We assessed ER stress upon exposure to FFAs by probing the cell death-inducing PERK/ATF4/CHOP arm of the UPR (Oyadomari and Mori 2004; Chan et al. 2015). Only treatment with EA or PA increased ATF4 and CHOP protein abundance, pointing to lipotoxicity-induced activation of the UPR (Fig. 4C). An increase in CPT1A protein abundance across all conditions showed successful intracellular delivery of all FFAs (Fig. 4C) (Thumelin et al. 1997). In line with previous work demonstrating that depletion of ER calcium is associated with lipotoxicity-induced ER stress (Oyadomari and Mori 2004; Marmugi et al. 2016), we observed significant reduction in ER Ca^2+^ levels only in cells treated with PA or EA (Fig. 4D). In contrast, cells treated with OA and PSA remained near-baseline, and cells treated with AA and GLA had increased ER Ca^2+^ levels (Fig. 4D).

Lipotoxic inflammation and recruitment of immune cells into pancreatic islets plays a central role in beta cell dysfunction and death in the context of T2D (Hotamisligil 2017). Based on the transcriptomic analysis (Fig. 2A) and previous work with PA (Eguchi et al. 2012), we focused on inflammatory signaling through NFkB measured by nuclear translocation of RELA (p65), a major component of NFkB-mediated transcription (Hayden and Ghosh 2008). Robust RELA nuclear translocation was noted after treatment with EA (Fig. 4E, S3B) and PA (Fig. 4E). In line with the transcriptomic analysis (Fig. 2A), PSA and OA triggered only modest RELA translocation (Fig. 4E) without affecting cell viability (Fig. 4A).

Next, we measured glucose stimulated insulin secretion (GSIS), a function specific to beta cells. Exposure to EA caused significant GSIS impairment (Fig. 4F). PA had a similarly deleterious effect (Fig. 4F)(Oh et al. 2018). In contrast, OA, PSA, AA and GLA had no effect on GSIS (Fig. 4F). Since impaired insulin secretion has been linked to disrupted autophagy (Ebato et al. 2008), and lipotoxicity increases autophagosome number (Mir et al. 2015), we quantified LC3B-positive autophagosomes in cells exposed to FFAs. Similar to PA, EA increased autophagosome numbers, whereas OA, PSA, AA and GLA had no effect (Fig. 4G-H). In sum, despite structural similarities with other MUFAs or very long chain FFAs, EA functionally behaved like the lipotoxic saturated FFAs (as summarized in Fig. 2D). Overall, these results reinforced the FFA clustering derived from systematic analyses, assigned lipotoxic effects to EA, and more broadly, to a previously unrecognized subset of 12 c2 MUFAs (Fig. 2D).

### c2 MUFAs are toxic to human cells

We took two additional approaches to assess the effects of toxic MUFAs such as EA in human pancreatic beta cells. First, taking advantage of advances enabling the differentiation of human induced pluripotent stem cells (iPSCs) into pancreatic beta cells (Pagliuca et al. 2014; K. Yang et al. 2020), we generated human iPSC-derived beta cells (Maxwell and Millman 2021; Hogrebe et al. 2020), and treated them with either EA (putative toxic MUFA) or OA (putative benign MUFA). Exposure to EA induced cell death in a dose and time-dependent manner, whereas OA had no effect on cell viability (Fig. 4I,J). In a complementary approach, we studied human pancreatic islets acutely isolated from cadaveric donors. We treated these islets with MUFAs representing all three MUFA-containing clusters c2, c3, and c4 (Fig. 4K,L). Similar to MIN6 cells and iPSC-derived human beta cells, c2 MUFAs (7 FFAs including EA) were toxic to human islet beta cells in a dose-dependent manner, whereas c3 (oleic and petroselenic acid) and c4 MUFAs (10(Z)-nonadecenoic acid) had no effect on cell viability (Fig. 4K,L). We concluded that the c2 MUFAs can induce significant injury, making them highly relevant to human beta cell biology.

### Structural characterization of c2 MUFAs

After confirming the toxicity of c2 MUFAs in both mouse and human beta cells, we next asked whether molecular or chemical features could predict and explain this newly defined FFA grouping. Recognizing that double bond number alone did not account for transcriptome-based clustering, we generated a matrix of all 2D FFA structural features (Methods). A random forest classifier trained on the pre-processed structural feature matrix successfully predicted assignment to transcriptome-based predefined clusters c1, c2 and c5 with high sensitivity and specificity based on leave-one-out cross validation (Fig. S3C; Methods). This classifier showed that while the number of double bonds was an important distinguishing feature (Fig. 5A, b_double), the longest chain of single bonds (b_max1len) and bond rotation (b_rotR) were highly predictive for c2 cluster assignment even amongst the MUFAs (c1-c4; Fig. 5A). The longest chain of single bonds also reliably captured the single PUFA (13(Z), 16(Z), 19(Z)-docosatrienoic acid) that was transcriptomically assigned to c2, and predicted its separation from the rest of the PUFAs in the c5 cluster (Fig. 5A).

**Fig. 5.**
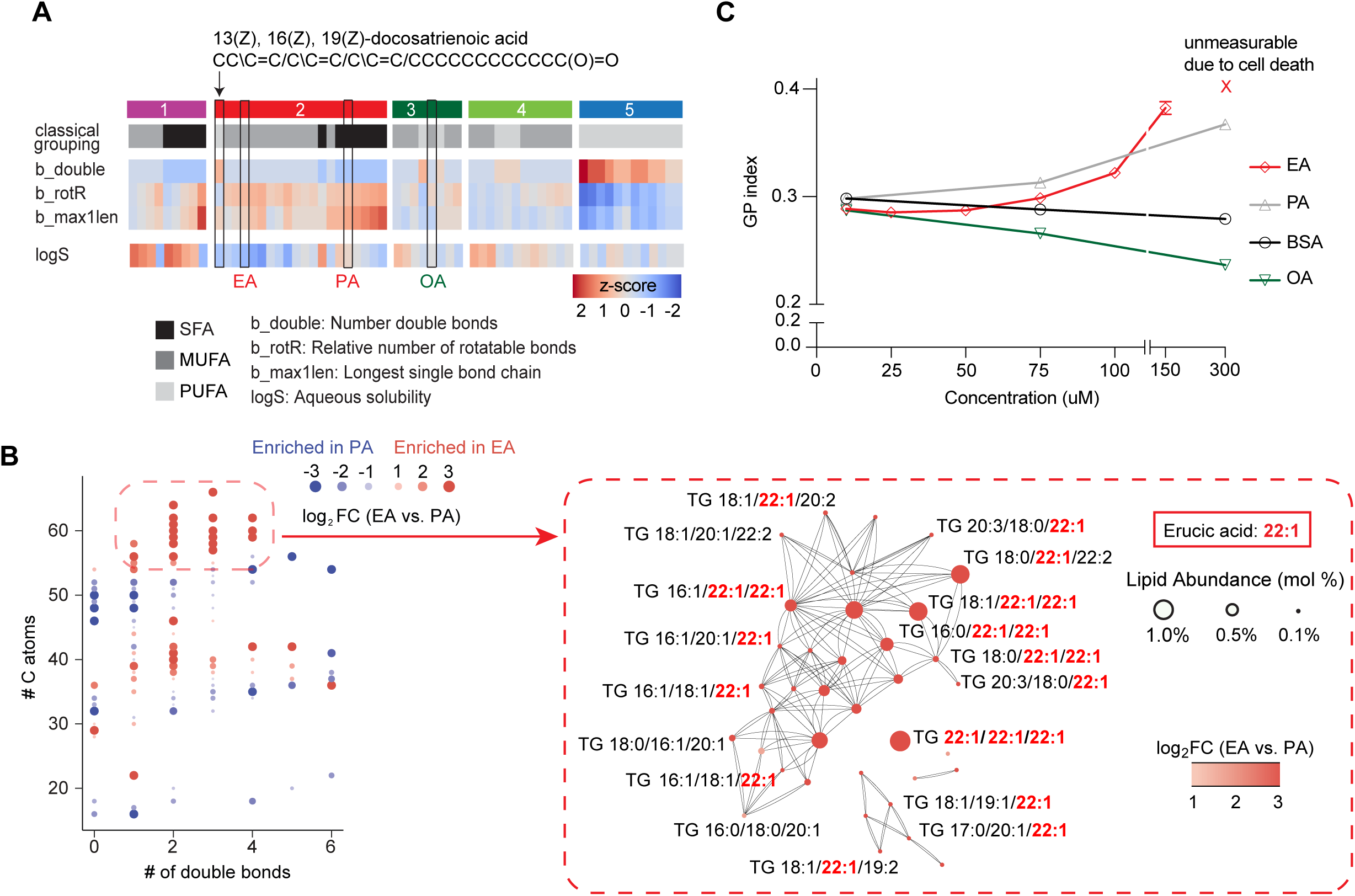
Long single bond chain in MUFA (EA) induces a distinct lipidomic profile that is associated with increased membrane rigidity. (A) Selected features from the decision tree analysis based on meta-features of highest importance (Mean Decrease Accuracy; methods). The longest single bond chain is the meta-feature that predicts the inclusion of 13(Z), 16(Z), 19(Z)-do- cosatrienoic acid as the only PUFA in the c2 cluster, and distinguishes between toxic EA and non-toxic OA. (B) Accumulation of longer unsaturated acyl chains found by lipidomic analysis of MIN6 cells treated for 24 h with 500 µM EA (left). A network analysis of the biochemical relationship (lines) between significantly enriched lipid species in the EA-induced lipidome highlighted the accumulation of EA (22:1)-containing triglyceride (TAG) species (right). (C) Membrane rigidity as measured by the GP index of Laurdan dye in INS1E beta cells after 12 h treatment with 3-6 differ- ent concentrations of BSA, PA, EA, or OA. PA increases membrane rigidity proportionally to its concentration; EA decreases membrane rigidity at low concentration similar to OA, but at higher, toxic levels, EA increases membrane rigidity similar to PA. Data are mean ± SEM; n=6 wells.

### Erucic acid induces a lipidome distinct from palmitic acid that is associated with changes in membrane fluidity

Exposure to PA leads to an increase in the abundance of saturated acyl chains of more complex lipid species (Piccolis et al. 2019). This in turn is linked to the activation of the UPR, which can sense changes in lipid composition or lipid bilayer stress (Olzmann and Carvalho 2018; Promlek et al. 2011; Volmer, van der Ploeg, and Ron 2013; Shen et al. 2017; Ho et al. 2020). Since c2 MUFAs caused significant cellular injury, similar to PA, we hypothesized that exposure to c2 MUFAs with long single-bond chains (such as EA) may induce specific changes to the cellular lipidome that could explain their lipotoxic effect. We performed a lipidomics analysis and found that exposure of cells to EA led to the accumulation of longer, unsaturated acyl chains in multiple lipid classes (DAGs, PCs, PEs), and especially in triglycerides (TAGs)(Fig. 5B, S3D). This profile was consistent with the integration of the very long-chain unsaturated EA into more complex lipids, and represented a lipidome distinct from that induced by PA (Fig. 5B, Table S2).

To probe the functional consequences of incorporation of long-chain FFAs into the lipidome, we measured membrane fluidity using a Laurdan dye that fluoresces at different wavelengths in accordance with lipid order/disorder, following a protocol previously used to demonstrate maladaptive PA-induced membrane rigidity (Golfetto, Hinde, and Gratton 2013; Pérez-Martí et al. 2022). Notably, EA had a unique biphasic effect on membrane fluidity. At the lowest concentrations, EA was similar to OA with a trend towards increasing membrane fluidity (Fig. 5C). At higher concentrations, similar to those measured in plasma from patients on FFA-rich diet (Aubourg et al. 1993; Lust et al. 2021) (and at which we found that EA is toxic (Fig. 4B)), membrane fluidity was decreased (Fig. 5C). Of interest, the membrane regions that were most rigid in EA treated cells differed from those in PA treated cells. Exposure to PA led to linear regions of increased rigidity reminiscent of the ER (Ruiz et al. 2019, 2021)(Fig. S3E). In contrast, EA induced distinct spherical regions of increased rigidity reminiscent of lipid droplet morphology (Fig. S3E). These observations are consistent with robust EA incorporation into TAGs (Fig. 5B). In sum, we found that exposure to EA generated a distinct cellular lipidome characterized by species with long unsaturated acyl chains that induced toxicity by decreasing cellular membrane fluidity.

### Identification of genes at the intersection of FFA exposure and genetic risk for metabolic disease

Complex diseases arise from the interaction of genetic risk and environmental exposures (Dempfle et al. 2008). GWAS studies have greatly contributed to our understanding of the genomic architecture of complex diseases (Claussnitzer et al. 2020), including T2D (Mahajan et al. 2014; Petrie, Pearson, and Sutherland 2011; Fuchsberger et al. 2016). While several T2D genomic studies have shown a strong association with loci linked to beta cell function (Pasquali et al. 2014; Flannick et al. 2014; Dwivedi et al. 2019; Thomsen et al. 2018), it remains challenging to annotate variants and prioritize genes for mechanistic functional follow-up studies (www.icda.bio).

In our experiments, beta cell exposure to 20 c2 FFAs significantly up- or down- regulated specific genes (Fig. 2A, Fig. S2A). We hypothesized that a subset of these genes could also be implicated in genetic risk for T2D. Based on this hypothesis, we sought to prioritize disease-relevant genes by investigating the overlap between an annotated large-scale T2D GWAS dataset (Mahajan et al. 2018) and genes highlighted by our transcriptomic analysis (Fig. 2A, Fig. S2A).

Taking advantage of the Multimarker Analysis of GenoMic Annotation tool (MAGMA; (de Leeuw et al. 2015); Methods), we generated a ranked list of T2D genes. We used the MAGMA gene set analysis (GSA) function (de Leeuw et al. 2015) to test whether the beta cell c2 FFA transcriptomic signature was enriched in the MAGMA- ranked T2D genes. Overlapping of the two datasets revealed that the top 5% of significantly differentially expressed lipotoxicity genes were enriched among the T2D GWAS ranked genes (FDR < 0.001, Fig. 6A). This enrichment was specific to T2D as compared to GWAS datasets for type 1 diabetes, an autoimmune disease (Onengut-Gumuscu et al. 2015) or schizophrenia, an unrelated neurodevelopmental disorder (Schizophrenia Working Group of the Psychiatric Genomics Consortium 2014); Fig. 6A). T2D GWAS genes were specifically enriched in the signature of the c2 cluster but not in the other four clusters (Fig. 6B). These results independently validated the utility and human relevance of FALCON for gaining new insights into FFA-associated metabolic diseases such as T2D (Fig. 1A, tier 2). Next, we plotted the genes that drove this enrichment according to their MAGMA rank on the x-axis (Table S3) and their lipotoxicity rank on the y-axis (based on differential expression p-value, Table S4, Fig. 6C). To highlight genes that drove the observed enrichment, we conservatively chose the top 5% lipotoxicity gene set as the y-axis boundary (464 genes) and the top 600 genes of the ranked MAGMA list as the x-axis boundary. We thus identified 25 genes at the intersection of lipotoxicity and T2D (Fig. 6C,D).

**Fig 6.**
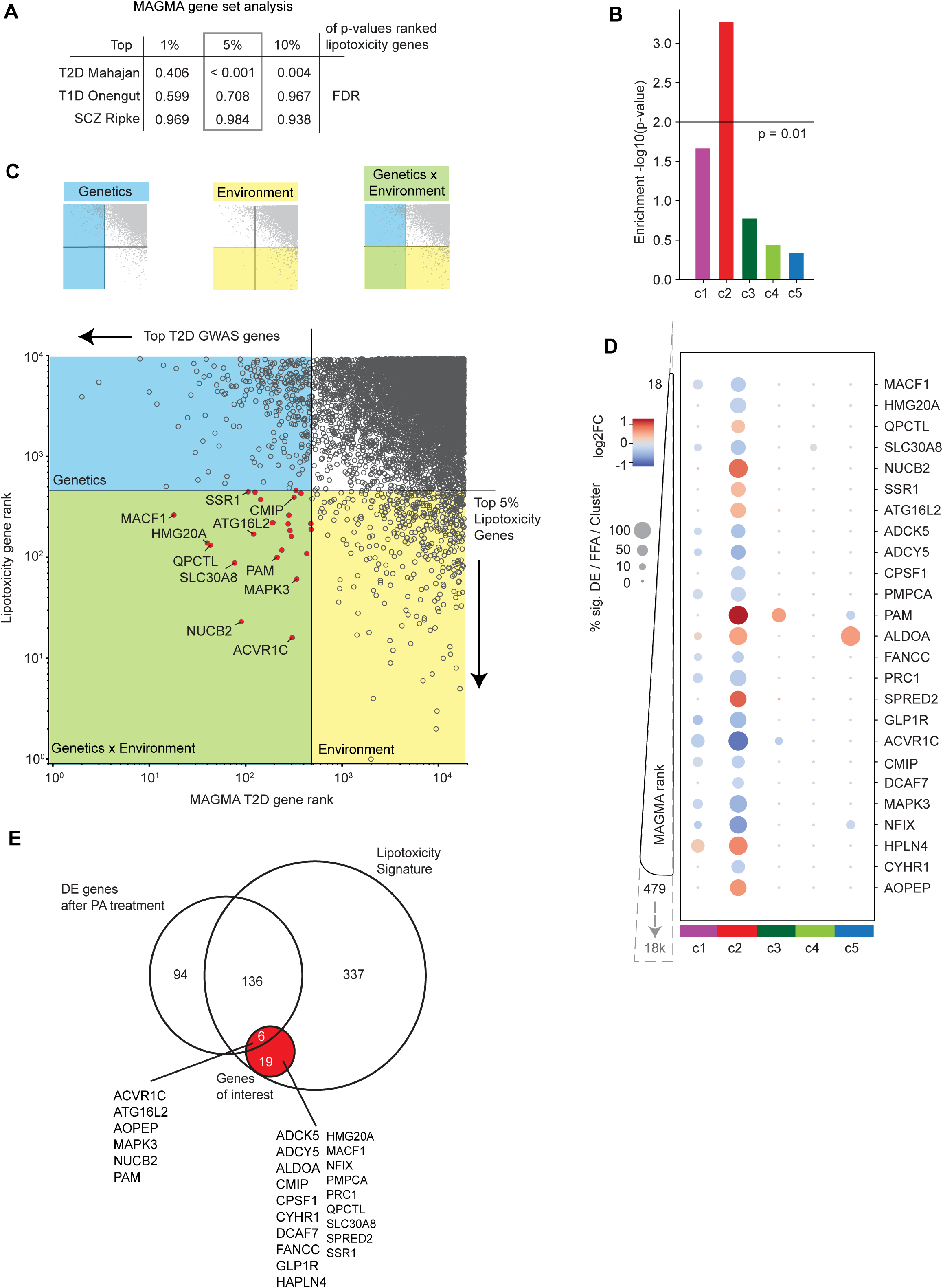
Integration of lipotoxicity transcriptomic signature with T2D GWAS dataset identifies *CMIP* as mediator of genetic and environmental risk for disease. (A) Gene set analysis (GSA) results based on the c2 lipotoxicity signature. T2D GWAS genes were ranked based on MAGMA score. A T1D and a schizophrenia GWAS dataset served as negative controls. Lipotoxicity gene sets are defined as top DE genes (1, 5, 10%) in transcriptomic profiles from the c2 cluster ranked by p-value and log2 fold change (LFC). GSA showed significant enrichment (FDR<0.05) for the top 5% (boxed) and 10% lipotoxicity gene sets. (B) Enrichment analysis for all FFA clusters (top 5% gene set) revealed that the c2 cluster gene set is uniquely enriched in the T2D GWAS dataset. (C) Scatter plot of genes based on T2D MAGMA rank (x-axis) and lipotoxicity rank (y-axis). Horizontal boundary defines top 5% lipotoxicity signature genes, vertical boundary defines top 600 MAGMA ranked T2D genes. As demonstrated in the schematic (top), genes located in the left (blue) quad- rants are associated with T2D genetic risk, genes located in the bottom (yellow) quadrants are associated with lipotoxicity environmental risk, and genes located in the bottom left (green) quad- rant are associated with both genetic and environmental risk. Genes of interest (red) drove the enrichment of the lipotoxicity signature in the GWAS dataset. (D) Expression pattern for the top 25 overlapping T2D-lipotoxicity genes across all FFA clusters. Size of dots represents the percentage of FFAs/cluster that induce significant differential expression of corresponding gene (p<0.05, Ben- jamini-Hochberg), colors represent strength and directionality of transcriptional changes (log2 fold change). (E) Venn diagram comparing the results of the analysis using the PA-induced signature alone compared with using the c2 lipotoxicity signature derived from all 20 lipotoxic FFAs. 19/25 genes, including *GLP1R* and *CMIP*, would have been missed if the analysis was limited to the PA-induced signature alone.

The sensitivity by which we could detect genes of interest was greatly enhanced by the fact that the lipotoxicity signature used in our analysis was based on an integrated transcriptome derived from all 20 lipotoxic c2 FFAs (including the newly defined lipotoxic MUFAs). Accordingly, the transcriptomic signature generated by PA treatment alone could predict only 6 of the 25 T2D-lipotoxicity genes, thus missing 19 important gene candidates, among them *GLP1R*, *SLC30A8, ADCY5,* and *CMIP* (Fig. 6E). GLP1R is a G-protein coupled receptor, the drug target for a class of medicines known as incretin mimetics (GLP1R agonists) approved for the management of obesity and T2D (Winzell and Ahrén 2008; Doyle-Delgado et al. 2020; Blonde et al. 2006; Wadden et al. 2013; Wilding et al. 2021). *SLC30A8* encodes the zinc transporter ZnT8 which plays an important role in insulin storage and glucose tolerance (Lemaire et al. 2009; Pound et al. 2009; Nicolson et al. 2009; Wijesekara et al. 2010). Variants in *SLC30A8* modify T2D risk (Flannick et al. 2014; Dwivedi et al. 2019). T2D risk alleles are also associated with decreased *ADCY5* expression in human islets (Roman et al. 2017). The identification of biologically and therapeutically relevant targets such as *GLP1R* directly validated our approach of using an integrated lipotoxicity signature derived from all 20 c2 FFAs to identify genes of interest at the intersection of genetic and lipotoxic risk for disease. Of the T2D-lipotoxicity genes revealed by our analysis, we focused our next set of experiments on *CMIP* because (i) it had not been previously implicated in beta cell biology (despite its poorly understood association with the *Maf* transcription factor family that bears connections to islet biology (Kataoka et al. 2004)) and (ii) it had no known role in lipotoxicity, thus offering opportunities for new biological insights.

### CMIP suppresses lipotoxicity in beta cells

C-MAF inducing protein (CMIP) has been implicated in kidney disease (Moktefi et al. 2016; Bouachi et al. 2018) and cancer (J. Zhang et al. 2017), but it has not been studied in beta cells. Of interest, CMIP-associated risk loci are linked to alterations in body mass index (Cao et al. 2018) and dyslipidemia (Mo et al. 2018). In immune cells, CMIP interacts with RELA and reduces NFkB activation (Kamal et al. 2009). A genome-wide interaction analysis with the insulin secretion locus MTNR1B identified an interaction with a CMIP intronic SNP affecting T2D risk (Keaton et al. 2018). To study *CMIP* in beta cells, we generated MIN6 *Cmip* knockout cell lines using CRISPR/Cas9, and isolated one *Cmip* knockout (CMIP KO) clone with a complete deletion of the major CMIP isoform (Fig. S4A). At baseline, this CMIP KO line and the non-targeting guide WT control displayed (i) similar morphology (Fig. S4B), (ii) expressed key beta cell markers (qPCR; Fig. S4C), (iii) responded to high glucose by a 2-fold increase in insulin secretion (Fig. S4D), and (iv) showed similar doubling rates in culture (Fig. 7A). These data suggested that at baseline CMIP is not required for beta cell survival or insulin secretion. In contrast, *CMIP* deletion significantly increased sensitivity to lipotoxic stress and cell death (Fig. 7B,C). Specifically, *CMIP* deletion exacerbated the lipotoxic effects of EA, and it even converted the typically non-toxic c3 MUFAs oleic and petroselenic acid into toxic FFAs (Fig. 7B). Functionally, *CMIP* deletion increased NFkB signaling (Fig. 7D,E)(Kamal et al. 2009). In CMIP KO cells, EA shifted the peak of RELA nuclear translocation from 18 hours to 3 hours while increasing the overall peak and baseline signal (Fig. 7D). Exposure to PA or EA, but not PSA or OA, increased RELA nuclear translocation in CMIP KO cells more than 4-fold compared to WT controls (Fig. 7E). Thus, CMIP deletion specifically increased inflammatory signaling in beta cells in response to lipotoxic FFAs. CMIP deletion also worsened the EA- or PA-induced reduction in insulin secretion (Fig. 7F). Overall, CMIP deletion rendered beta cells more vulnerable to FFA exposure.

**Fig 7.**
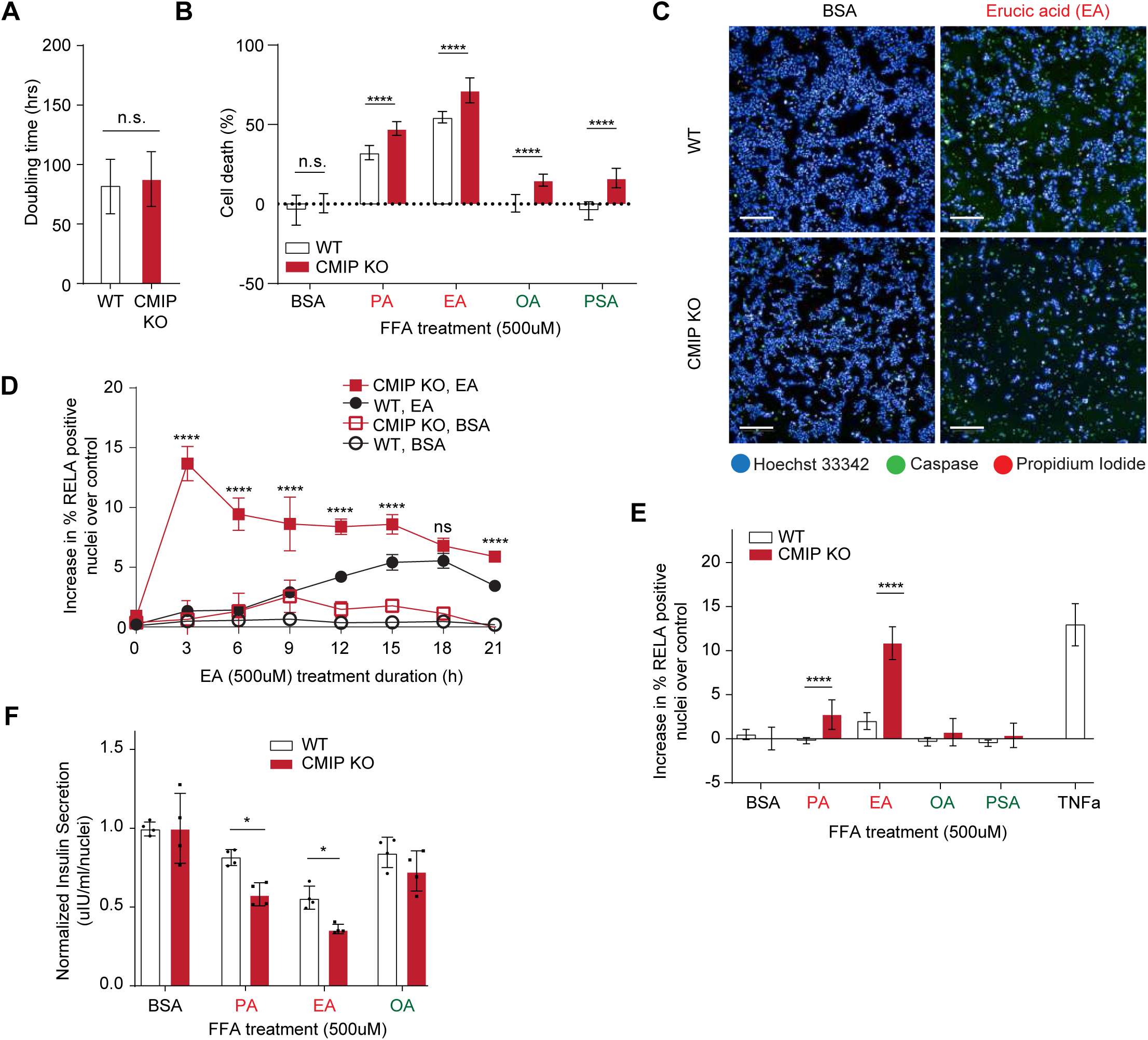
CMIP deletion sensitizes beta cells to FFA-mediated injury and cell death. (A) CMIP deletion does not affect cell proliferation. No difference in the doubling rate between WT and CMIP KO cells (Two-sided t-test, n=19 splits). (B) After exposure to lipotoxic PA or EA, cell death is increased in CMIP KO compared to WT cells. Non-toxic OA and PSA are rendered toxic in CMIP KO cells. Cell death was measured as number of viable (caspase 3/7 and propidium iodide negative) cells compared to the non-treated control for WT and CMIP KO cells after 72 h exposure to FFAs (500 μM, n=21 wells). Data are mean ± SD. Two-way ANOVA with multiple comparisons (Šídák correction,****p < 0.0001). (C) Representative images of the cell death assay showing increased susceptibility to EA in CMIP-deleted cells. Nuclei were marked by Hoechst and apoptosis was measured by a caspase 3/7 dye (green). Dead cells were stained with propidium iodide (red). Scale bars: 100 μm. (D) Percentage of cells with RELA nuclear translocation after exposure to EA or BSA (500 μM) at 3 h intervals from 0 to 21 h after subtrac- tion of baseline signal from non-treated cells. Data are mean ± SD, n = 7 wells. Two-way ANOVA with multiple comparisons (Šídák correction,****p < 0.0001). (E) Percentage of cells with RELA nuclear translocation after 3 h exposure to FFAs (500 μM) or TNFα (50 ng/ml; positive control) after subtraction of baseline signal from non-treated cells. CMIP KO increased the percentage of RELA translocation after PA or EA treatment. Data are mean ± SD. Two-way ANOVA with multi- ple comparisons (Šídák correction, ****p < 0.0001). (F) Insulin secretion at baseline normalized to BSA control after 24 h treatment with FFA (500 μM). Insulin secretion was reduced in CMIP deleted cells upon exposure to lipotoxic FFAs. Data are mean ± SD. Two-way ANOVA with multi- ple comparisons (Holm-Šídák correction, *p < 0.05). PA, palmitic acid; EA, erucic acid; PSA, petroselenic acid; OA, oleic acid.

To confirm that the observed phenotypes could be attributed to the loss of CMIP, we next tested whether reintroducing CMIP in CMIP KO cells could reverse its deleterious effects. Restoring CMIP abundance in CMIP KO cells (as quantified in Fig. S4F) led to a partial rescue from EA-induced cell death (Fig. 8A) and to a faster decline in EA-induced inflammatory NFkB signaling (Fig. 8B; no effect on NFkB signaling in the absence of EA (Fig. S4G)). Similarly, glucose stimulated insulin release was improved upon restoring CMIP expression, especially in EA-treated cells (Fig. 8C). Taken together, these data revealed that CMIP, initially found among thousands of loci in a T2D GWAS, could now be prioritized as a putative suppressor of lipotoxic injury in beta cells.

**Fig 8.**
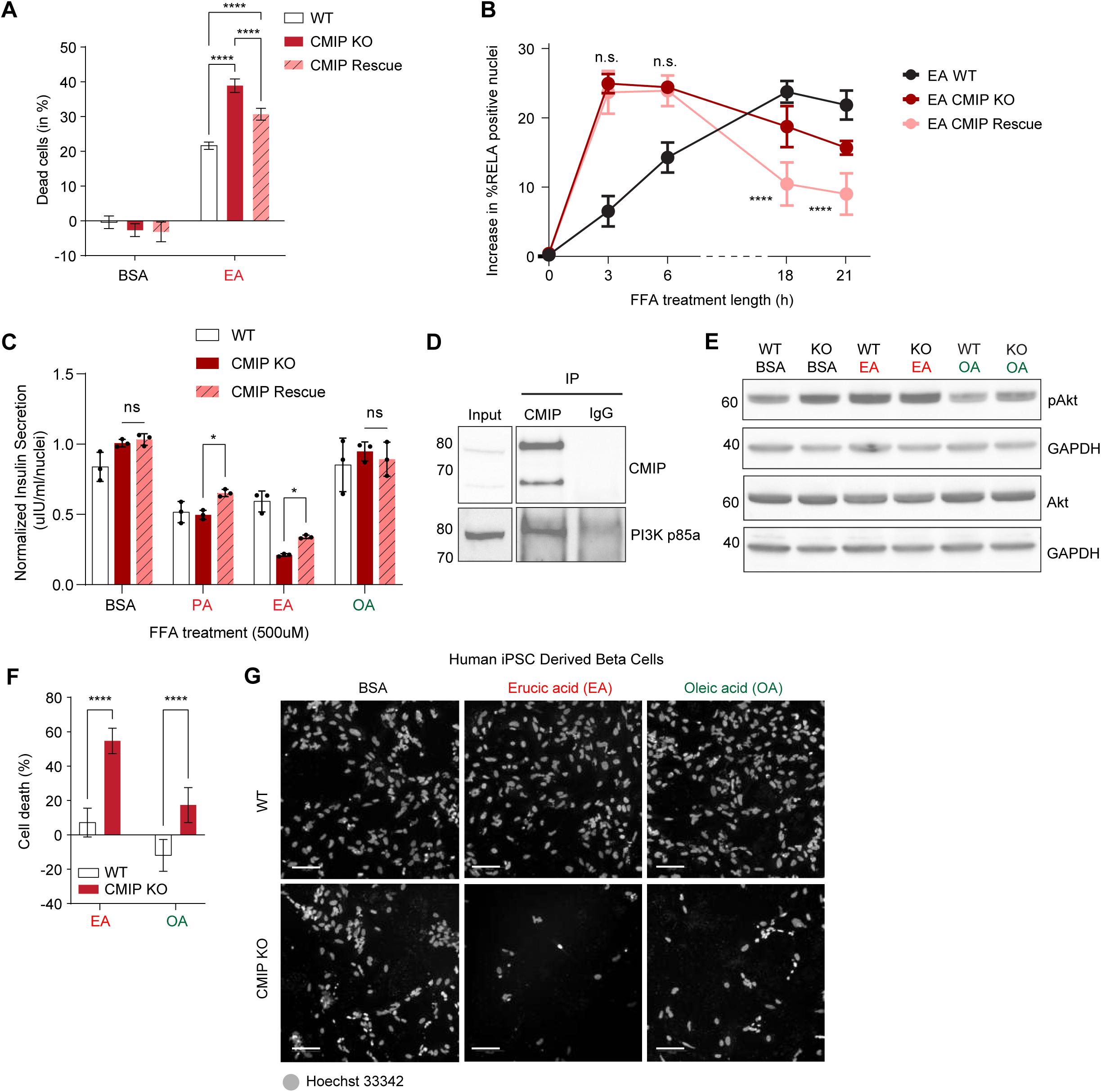
CMIP protects beta cells from FFAs by regulating Akt activity, and human beta cells lacking CMIP are vulnerable to FFA-induced cell death. (A) Reintroduction of CMIP in CMIP deleted cells (CMIP rescue) attenuates the toxic effects of EA. Percentage of cell death as measured by decreases in MIN6 viable cells compared to the non-treated control after 24 h exposure to FFAs (250 μM, n=9 wells). Data are mean ± SD. Two-way ANOVA with multiple comparisons (Tukey correction,****p < 0.0001). (B) CMIP rescue attenuates inflammatory NFkB signaling. Percentage of cells with RELA nuclear translocation upon treatment with EA (500 μM) for the indicated times after subtraction of baseline signal from non-treated cells. Data are mean ± SD, n = 6-9 wells. (C) CMIP rescue partially restores insulin secretion in cells exposed to c2 FFAs. Insulin secretion upon glucose stimulation normalized to BSA control for WT and CMIP KO cells after 24 h treatment with FFA (500 μM). Data are mean ± SD. Two way ANOVA with Šídák multiple comparison test (*p < 0.05, n=3 wells). (D) PI3K p85ɑ immunoprecipitates with CMIP in beta cells. Western blot displaying lysate input (left) or co-IP with a CMIP or IgG control antibody (right) stained for CMIP (top) or PI3K (bottom) (n=3 blots). (E) Phosphorylated Akt (pAkt) abundance is increased in CMIP KO cells compared to WT controls. EA exposure increases pAkt in WT cells, but not in CMIP KO cells indicating that CMIP deletion maximizes pAkt levels at baseline. Western blot for pAkt and total Akt after 24 h treatment with 500 μM FFAs (GAPDH, loading control; n=3 blots). (F) In human iPSC-derived beta cells, CMIP deletion promotes cell death, in agreement with experiments in MIN6 beta cells (Fig. 7B). Cell death in human iPSC-derived beta cells as measured by decreases in cell count compared to BSA-treat- ed control. Cells were treated with BSA, EA, or OA at 500 µM for 24 h (n=24 wells). (****p<0.0001, two-way ANOVA with Bonferroni multiple comparison test). (G) Representative images of cell death assay in human iPSC-derived beta cells as measured by number of nuclei (Hoechst). Scale bars: 100 μm. PA, palmitic acid; EA, erucic acid; OA, oleic acid.

To gain insights into CMIP’s function in beta cells, we analyzed transcriptomic data from human islets (Taneera et al. 2012). *CMIP* gene expression correlated with several pathways including PI3K-Akt signaling, insulin secretion, FFA metabolism, and AMPK signaling (Fig. S4H,I), suggesting that CMIP may play a role in metabolic regulation. Intriguingly, consistent with prior work in peripheral blood mononuclear cells (Kamal et al. 2010), we found that that the regulatory subunit of PI3K (p85L) co-immunoprecipitated with CMIP in beta cells (Fig. 8D), suggesting that this interaction may connect CMIP to metabolic signaling pathways (Fig. S4H,I).

To experimentally probe these pathways, we assessed the activity of three critical metabolic sensors in beta cells: AMPKL signaling were unchanged after CMIP deletion (Fig. S4J). In contrast, CMIP deletion resulted in increased pAkt protein abundance (with no effect on total Akt, Fig. 8E). Interestingly, pAkt in WT cells exposed to EA was at the same level as pAkt in CMIP KO cells at baseline (Fig. 8E). When exposed to EA, CMIP KO cells could not increase pAkt further (Fig. 8E). Since modulation of Akt signaling in response to metabolic stress is thought to promote beta cell survival (Camaya, Donnelly, and O’Brien 2022; Elghazi and Bernal-Mizrachi 2009), and PI3K activity increases the presence of phosphoinositide signaling molecules (PIP2 and PIP3) in the plasma membrane to recruit Akt for activation and downstream signaling (Long et al. 2021), the simplest explanation for our data is that CMIP modulates Akt activity through its interaction with PI3K. Upon CMIP deletion, cells lose the ability to dynamically regulate Akt, making them more vulnerable to FFA-induced cell death compared to WT beta cells (Fig. 7B).

As a final test, and to probe for the human relevance of CMIP, we generated human iPSC-derived beta cells in which *CMIP* was deleted. In these human cells, *CMIP* deletion reduced cell viability after treatment with either EA or OA (Fig. 8F,G). In sum, exposure to excess FFAs was particularly deleterious to human beta cells lacking CMIP, validating the key role of CMIP in a relevant model for human health and disease.

## Discussion

Lipids, including FFAs, are ubiquitous in living organisms and essential for life, and yet, significant knowledge gaps remain in our understanding of FFA biology. In this study we pioneered FALCON, a multimodal, systematic approach to functionally characterize structurally diverse FFAs. Our approach was comprehensive both in input (number of FFAs tested) and output (multimodal readouts). The added dimensionality - a direct consequence of studying the effects of 61 diverse FFAs - provided the necessary power to uncover biological features across the entire spectrum of FFAs tested. Our studies led to several important conclusions.

First, 20 structurally diverse FFAs defined the toxic (c2) cluster, a group of FFAs united solely by the fact that they mediated similar functional outcomes. The identification of MUFAs in the c2 cluster (12/20 FFAs) suggests a shift in how we interpret the toxicity of FFAs, because we show that saturation alone is not sufficient to predict the lipotoxic potential of a given FFA. MUFAs like OA have been proposed to have harmless or even beneficial effects (Palomer et al. 2018). However, our experiments, including studies in human islets, kidney epithelial cells and microglia showed that OA is not representative of the entire MUFA class, and highlighted several MUFAs, such as EA, that were highly toxic. Our classifier (Fig. S3C) showed that the longest chain of single bonds was predictive of c2 MUFA toxicity, correlating with toxic SFAs rather than non-toxic MUFAs in other clusters (c3 and c4). These findings suggest that a double-bond containing MUFA with a long stretch of carbons linked by single bonds is functionally similar to SFAs, at least in terms of cellular toxicity. Despite induction of similar transcriptomic responses, EA induced a lipidome that was distinct from that induced by PA. In contrast to the well-known accumulation of saturated acyl chains after PA treatment (Piccolis et al. 2019), the EA lipidome was characterized by an accumulation of long-chain unsaturated acyl chains. The distinct lipidomic profiles induced by EA versus PA prompted us to investigate why exposure to these FFAs leads to similar biological outcomes in beta cells (Fig. 4). The membrane fluidity studies hint at the mechanism behind this unique MUFA toxicity; at low concentrations, EA behaves like unsaturated FFAs (e.g. OA), but at higher concentrations, EA becomes harmful because the incorporation of its long unsaturated chains into complex lipid species changes the properties of lipid membranes in a manner similar to SFAs (e.g. PA). The resulting increase in membrane rigidity likely contributes to lipid bilayer stress culminating in activation of cell death pathways (Volmer, van der Ploeg, and Ron 2013). Future work may test whether increased membrane rigidity at high concentrations of EA is directly related to the predicted impact of incorporating its long chain of single bonds (Fig. S3C) into membrane lipids. In sum, we identified and characterized the biological effects of a previously unrecognized set of toxic MUFAs, a discovery enabled by our FALCON platform, and provided mechanistic insight into their particular behavior compared to toxic but structurally dissimilar SFAs.

Second, the comprehensive interrogation of many FFAs with FALCON allowed us to gain new biological insights. Due to the large number of FFAs studied simultaneously (e.g. 20 c2 toxic FFAs), we gained power well beyond that achievable by interrogating the effects of a single FFA alone (such as PA, Fig. 6E). Accordingly, we identified a set of 25 genes that are transcriptionally responsive to lipotoxic stress and are also associated with variants that confer genetic risk for T2D. Our analysis was internally validated by the identification of *GLP1R,* a well-known obesity (Wilding et al. 2021) and T2D drug target (Scott et al. 2016), and *SLC30A8*, a gene in which coding variants have been shown to modify risk for T2D (Krentz and Gloyn 2020; Flannick et al. 2014; Dwivedi et al. 2019). Importantly, we identified CMIP as a previously unrecognized suppressor of lipotoxicity, and we confirmed that these findings are relevant to humans using iPSC-derived beta cells. Future studies will focus on the precise molecular mechanism by which CMIP senses FFAs and modulates Akt and related pathways. Nevertheless, this proof-of-concept study illustrates FALCON’s ability to prioritize genes of high mechanistic value that may have otherwise gone unnoticed based on genomics data alone, and offers a method for prioritizing targets that reflect the combined effects of environmental exposure and genetic risk for disease.

Third, FALCON can serve as a valuable tool for exploring fundamental lipid biology in different cell types and tissues. Lipotoxicity and alterations in lipid metabolism have been implicated in numerous disorders including kidney disease (Morgado-Pascual and Opazo-Ríos 2020; Sidhom et al. 2021), neuropathy (Durán et al. 2019), cancer (Munir et al. 2019; Ringel et al. 2020; Ubellacker et al. 2020), liver disease (Liangpunsakul and Chalasani 2019), and Alzheimer’s disease (de la Monte and Tong 2014; Cutuli et al. 2020; Desale and Chinnathambi 2020; Chausse et al. 2019; Madore et al. 2020; Snowden et al. 2017). However, many of the fundamental mechanisms involving lipotoxic FFAs in these diseases have yet to be fully elucidated (Yoon et al. 2021). For example, metabolic alterations in cancer cells that increase fatty acid synthesis and uptake have long been correlated with cancer progression (Munir et al. 2019; German et al. 2016). In other tissues, such as the heart, studies exploring the mechanisms underlying lipotoxic cardiomyopathy have explicitly called for investigations into the contributions of individual fatty acids to disease pathogenesis (Nakamura et al. 2019). FALCON provides the fatty acid level resolution necessary to begin to tackle these complex questions across many cell types and diseases of interest.

Finally, an important translational implication of our study is the notion that the precise FFA profiles in human blood or tissue may carry valuable information about personalized disease risk and progression. Although extensive work is required to fully assess the implications of these findings *in vivo*, our results motivate the measurement of FFA species in serum samples from large cohorts of well-phenotyped patients. These patient-derived FFA profiles may generate a nuanced view of FFAs as an environmental risk factor, and integrate with the mechanistic insights generated from FALCON. We speculate that integration of polygenic risk scores for complex metabolic diseases (such as obesity, T2D and NAFLD) with serum or tissue-derived “lipotoxicity risk scores’’ for individual patients could become a useful tool for patient risk assessment and stratification in clinical trials.

## Limitations of the Study

Since some FFAs in our platform are protective while others are harmful, the study of FFA combinations may be of interest. We note that the large number of potential FFA combinations (>1800) currently limits the feasibility of such a systematic study, but future work may prioritize some FFA combinations for follow-up work.

While the work described in this manuscript was largely focused on c2 FFAs, we uncovered many additional biological processes that merit further study, for example, the putative role of ferroptosis induced by c5 PUFAs in microglia. By making all of our datasets publicly available, we hope that many colleagues in the scientific community will be empowered to explore them to gain fundamental insights into FFA biology in different cell types.

## Acknowledgements

We thank Katie Liguori for excellent graphic design work. We are also grateful to colleagues in the Center for the Development of Therapeutics for their assistance in the preparation of the FFA library. This work was supported by a seed grant from the Broad Institute of MIT and Harvard (B*n*10 to AG), who is also supported by DK095045, DK099465, and Cure Alzheimer’s Fund. NW was supported by a Deutsche Forschungsgesellschaft Fellowship (WI 4612/1-1) and is a participant in the BIH-Charité Junior Digital Clinician Scientist Program funded by the Charité –Universitätsmedizin Berlin and the Berlin Institute of Health, JCF was supported by the National Institute of General Medical Sciences (T32GM007753, T32GM007726), E-H.S. was supported by the National Institutes of Health (F30DK112477), C.K. was supported by the National Institutes of Health (F31DK126252), J.R. was supported by the National Institute of General Medical Sciences (T32GM007753), M.R.B was supported by the National Institutes of Health (K00DK123834), M.D.-L. was supported by a Broad–Israeli Science Foundation Fellowship, E.S. was supported by NIH 1-R01-DK126855-01 and an American Surgical Association Foundation Fellowship, MC was supported by FNIH AMP-T2D RFB8b, NIDDK UM1 DK126185, AEC and SS were supported by the National Institutes of Health (NIH MIRA R35 GM122547 to AEC), and RD was supported by Knut and Alice Wallenberg Foundation, Sweden (KAW 2019.0580). The content is solely the responsibility of the authors and does not necessarily represent the official views of the National Institutes of Health.

## Author Contributions

NW, JCF, JLP and AG conceived the study and designed the experiments. NW and JCF conducted experiments and analyzed the data; experimental work was contributed by CK, EHS, JLM, MDL, JSi, JP, JLP, JR, MT, HY, ZWL, AB, RD, JSm, VS, MH, and DL; data analysis was contributed by CK, EHS, MRB, CA, MKA, KRG, CC, HSA, SS, RS, AC, JF, and JLP. NW, JCF, JLP, JLS and AG wrote the manuscript; AG directed the study. All authors read the manuscript and approved its contents.

## Materials and Methods

### Cell Culture

MIN6 cells were purchased (Addex Bio, #C0018008) and cultured as described previously (Burns et al. 2015). In short, cells were maintained in DMEM with 4.5 g/L glucose, supplemented with 10% FBS (fetal bovine serum, Life Technologies, #26140079), 100 U/ml penicillin and 100 μg/ml streptomycin (#15140 Invitrogen) and 55 μM beta-mercaptoethanol (Sigma, #M6250). MIN6 cells were cultured and used for experiments up to passage 30.

Immortalized human kidney epithelial cells used in this study were maintained at 37°C with 5% CO2 in RenaLife Renal Basal Medium supplemented with RenaLife LifeFactors® (Lifeline Cell Technology), with the exclusion of gentamicin and amphotericin B. The cells were generated with informed consent under WFUHS IRB00014033.

INS-1E cells were grown in RPMI 1640 media, supplemented with 10% FBS, 1% penicillin and streptomycin, 1% sodium pyruvate, and 50 µM β-mercaptoethanol (all from Life Technologies). Cells were maintained in flasks pre-coated with supernatant from rat 804G cell line (804G matrix) as previously described (Parnaud et al. 2008).

Cell lines were routinely checked and were negative for mycoplasma.

### FFA Preparation

Enzo SCREEN-WELL^®^ Fatty Acid library (#BML-2803-0100) containing FFAs dissolved in DMSO ([FFA]stock = 10 µM) was stored in glass vials at -20°C in the compound management facility of the Broad Institute. Template plates for High Throughput Screens were stored up to 4 weeks at 4°C. To prepare compound plates, small volumes of DMSO dissolved FFAs were transferred into microplates containing fatty acid free BSA (Sigma #A8806) solutions in ddH2O in a molecular ratio of 1:6.67 (BSA:FFA, [FFA]final = 500 µM) with an automated simultaneous pipettor (Analytik Jena CyBio^®^ Well Vario). Plates were incubated overnight for 24 h at 37 °C to ensure complete binding of FFAs to BSA. Next, DMSO and ddH2O were completely removed with the GeneVac HT-12 evaporator for 12 h with full vacuum at 37 °C and continuous centrifugation at 400g. Plates with dry FFA bound BSA crystals in the wells were resuspended in MIN6 culture medium at room temperature for 4-8 h on an orbital plate shaker. After resuspension, compound plates were spun down at 5000g for 10 min and manually transferred to 384 MultiScreenHTS HV Filter Plates (0.45 µm, Millipore, #MZHVN0W10) and spun down again for 1 min at 500g into an empty compound plate. Resulting filtered compound plates were transferred into assay plates of the same format as the CyBio^®^ Well Vario simultaneous pipettor. Representative FFAs were ordered from Nu-Chek Prep, Inc., manually dissolved in DMSO ([FFA]stock = 10 µM) and prepared in glass vials according to the same protocol.

### Differential Scanning Calorimetry

All differential scanning calorimetry (DSC) measurements were performed with a MicroCal VP-Capillary DSC Automated system (Malvern Panalytical). Selected FFAs were bound to BSA in microplates according to the protocol described above and resuspended in PBS to a final concentration of [BSA]final = 50 µM. Sample measurements included one measurement of PBS vs. PBS to record a baseline reference curve at the scan rate of 200 °C/h. The samples were heated from Tstart = 10 °C up to Tend = 90 °C at the same scan rate. The melting temperature Tm was determined from the resulting single-peak melting curve using FFA-free BSA as a control.

### Lipid Profiling

40,000 MIN6 cells/well were seeded in 96 well plates 24 h prior to treatment in three replicates. FFA library compound plates were transferred into assay plates which were then incubated for 24 h. The lipid fraction of cells was isolated with isopropanol after washing the plates 3 times with ice cold PBS. After the addition of isopropanol, plates were incubated for 1 h at 4 °C. IPA extracts were then manually transferred to autosampler vials (Waters), capped, and stored at -80 °C until analysis. Lipid profiling was done as previously described (Paynter et al. 2018). Briefly, non-targeted liquid chromatography mass spectrometry (LC-MS) data were acquired using a system composed of a Nexera X2 U-HPLC system (Shimadzu Scientific Instruments; Marlborough, MA) coupled to a Q Exactive Focus Orbitrap mass spectrometer (Thermo Fisher Scientific; Waltham, MA). Cell extracts (2 µL) were injected directly onto a 100 x 2.1 mm, 1.7 µm ACQUITY BEH C8 column (Waters; Milford, MA). The column was then eluted isocratically with 80% mobile phase A (95/5/0.1; vol/vol/vol 10mM ammonium acetate/methanol/formic acid) for 1 min followed by a linear gradient to 80% mobile-phase B (99.9/0.1; vol/vol methanol/formic acid) over 2 min, a linear gradient to 100% mobile phase B over 7 min, then 3 min at 100% mobile-phase B. Mass spectrometry analyses were carried out using ESI in the positive ion mode using full scan analysis over 220–1100 m/z. Raw data were processed and visually inspected using TraceFinder 3.3 software (Thermo Fisher Scientific; Waltham, MA) and Progenesis QI (Nonlinear Dynamics; Newcastle upon Tyne, UK). The identity of individual metabolites and lipid families was confirmed by matching their retention time to that of authentic reference standards.

For the comparison between the PA and EA-induced lipidomes, MIN6 were seeded at 800,000 per well in a 6-well plate and grown for 3 days (n=4). After the cells were treated with the respective FFAs at 500 µM for 24h, cells were washed 3x with 1ml of room temperature PBS. Cells were scraped in 500ul room temperature PBS, pelleted at 13,000xg, and flash frozen in liquid nitrogen for storage after the supernatant was removed. Cell pellets were lysed in double distilled water following three cycles of freeze-thawing in a water bath sonicator. Subsequently, lipids were extracted according to Folch’s Method. The organic phase of each sample, normalized by tissue weight, were then separated using ultra high performance liquid chromatography coupled to tandem mass spectrometry (UHPLC-MSMS) (Narváez-Rivas and Zhang 2016). UHPLC analysis was performed employing a C30 reverse-phase column (Thermo Acclaim C30, 2.1x 250 mm, 3 μM, operated at 55° C; Thermo Fisher Scientific) connected to a Dionex UltiMate 3000 UHPLC system and a Q-Exactive Orbitrap high resolution mass spectrometer (Thermo Fisher Scientific) equipped with a heated electrospray ionization (HESI-II) probe. Extracted lipid samples were dissolved in 2:1 methanol:chloroform (v/v) and 5 μ of each sample was analyzed separately using positive and negative ionization modes, respectively. Mobile phase A consisted of 60:40 water/acetonitrile (v/v), 10 mM ammonium formate and 0.1% formic acid, and mobile phase B consisted of 90:10 isopropanol/acetonitrile (v/v), 10 mM ammonium formate and 0.1% formic acid. Lipids were separated over a 90 min gradient; during 0–7 minutes, elution starts with 40% B and increases to 55%; from 7 to 8 min, increases to 65% B; from 8 to 12 min, elution is maintained with 65% B; from 12 to 30 min, increase to 70% B; from 30 to 31 min, increase to 88% B; from 31 to 51 min, increase to 95% B; from 51 to 53 min, increase to 100% B; during 53 to 73 min, 100% B is maintained; from 73 to 73.1 min, solvent B was decreased to 40% and then maintained for another 16.9 min for column re-equilibration. The flow-rate for chromatographic separation was set to 0.2 mL/min. The column oven temperature was set at 55° C, and the temperature of the autosampler tray was set to 4° C. The spray voltage was set to 4.2 kV, and the heated capillary and the HESI were held at 320° C and 300° C, respectively. The S-lens RF level was set to 50, and the sheath and auxiliary gas were set to 35 and 3 units, respectively. These conditions were held constant for both positive and negative ionization mode acquisitions. External mass calibration was performed using the standard calibration mixture every 7 days. MS spectra of lipids were acquired in full-scan/data-dependent MS2 mode. For the full-scan acquisition, the resolution was set to 70,000, the AGC target was 1e6, the maximum injection time was 50 msec, and the scan range was m/z = 133.4–2000. For data-dependent MS2, the top 10 precursor selection in each full scan were isolated with a 1.0 Da window, fragmentation using stepped normalized collision energy of 15, 25, and 35 units, and analyzed at a resolution of 17,500 with an AGC target of 2e5 and a maximum injection time of 100 msec. The underfill ratio was set to 0. The selection of the top 10 precursors was subject to isotopic exclusion with a dynamic exclusion window of 5.0 sec. All data were analyzed using the LipidSearch version 5.0 SP (Thermo Fisher Scientific) and all identified species (grade A, B) were reported.

### RNASeq / SmartSeq 2

40,000 MIN6 cells/well were seeded in 96-well plates 24 h prior to treatment in three replicates. FFA library compound plates were transferred into assay plates which were then incubated for 24 h (n=6). RNA was extracted from cells using TCL buffer (#1031576, Qiagen) with 1% beta-mercaptoethanol followed by an RNA clean-up with Agencourt RNA cleanup XP (#A63987, Beckman Coulter). Bulk RNA (1 μl) was added to a 3 step cDNA synthesis reaction with 3’RT (5’- AGCAGTGGTATCAACGCAGAGTAC(T30)VN-3’, IDT), template switching (5’- AAGCAGTGGTATCAACGCAGAGTACrGrG+G-3’, Qiagen), and ISPCR (5’-AAGCAGTGGTATCAACGCAGAGT-3’, IDT) oligos from the SMART-seq2 protocol (Picelli et al. 2014). cDNA was purified using AMPure XP Agencourt (#100609, Beckman Coulter) and quantified using Qubit dsDNA High Sensitivity (#102689, Life Technologies). Samples were diluted to 0.2 ng/ul in TE and tagmented (Nextera XT DNA Library Preparation Kit (#FC-131-1096, Illumina). Indexing was performed using the Nextera XT Index Kit (#FC-131-1001, Illumina). Final libraries were QCed using the Qubit dsDNA High Sensitivity kit and Bioanalyzer High Sensitivity DNA Kit (#5067-4627, Agilent). Libraries were sequenced at a concentration of 1.8 pM on a NextSeq with a 75 cycle v2 kit (#TG-160-2002, Illumina) with a read structure of Read 1 37bp, Read 2 37bp, Index 1 8bp, and Index 2 8bp. Each sample had approximately 4 million reads.

### Cell Painting

MIN6 cells were seeded at a density of 15,000 per well in a 384-well plate (Perkin Elmer, CellCarrier Ultra, #6057300). After 24 h recovery, cells were treated with the FFA library and incubated for 24 h. Cell staining and fixation was performed according to previous protocols (Bray et al. 2016). Images were acquired with the Opera Phenix High Content Screening System (#HH14000000, Perkin Elmer) with a 63X/1.15NA water immersion lens. Image quality control was carried out using CellProfiler 2.2.0 (Kamentsky et al. 2011) and CellProfiler Analyst (Dao et al. 2016) according to a prior machine-learning based protocol (Bray et al. 2012). Image illumination correction and analysis were performed in CellProfiler (pipelines available upon request). After analysis, the data were compiled and normalized in Cytominer (code available at https://github.com/cytomining/cytominer) as described previously (Bray et al. 2016). Briefly, single cell level features were aggregated per well by computing averages. The mean values and dispersion of the 2202 features measured in all samples were normalized to negative controls (BSA). Features with near-zero variance were removed, and a non-redundant feature set was created by inspecting pairwise correlations. The remaining 1222 features comprise the morphological profile of a given well.

### Data Analysis

Unless otherwise stated all computational and statistical analysis in this study has been performed in Python and R.

#### (1) Lipidomics

A blocked experimental design with one replicate of each FFA in the library, together with multiple BSA controls per 96-well plate, was chosen (n=3). Raw lipidomic profiles received from the Metabolomics Platform at the Broad Institute were filtered for samples with strongly deviating sample medians (manual cutoff, 7 out of 280 or 3% of the samples were discarded). Lipid metabolites with more than 30% of missing data points were removed, otherwise missing values were substituted with 50% of the minimum value of the respective metabolite’s intensity. To account for variations in total amount of captured metabolites, samples were scaled towards the global sample median. Only annotated lipid metabolites were used for further differential abundance analysis. We sought to understand the relationship between structural features of externally added FFAs and changes in the triglyceride fraction of the cells (Fig. 1C). For each externally added FFA, triglyceride intensity deviations from the BSA control were summed based on the structural feature of interest (number of C-atoms, number of double bonds). Then, triglyceride profiles of externally added FFAs were summarized based on the structural feature of interest of the FFAs (number of C-atoms, number of double bonds) and normalized to the number of FFAs making up each group. For assessment of the global lipidome in response to erucic and palmitic acid, lipid metabolites were filtered as described and subsequently imported into lipidR (Mohamed, Molendijk, and Hill 2020).

Differential analysis of lipid abundance was calculated using the empirical Bayes procedure. Fold change in lipid abundance (EA vs. PA) was then normalized based on noted structural features (number of C-atoms, number of double bonds) and visualized in R. Network analysis of the biochemical relationship between differentially abundant lipid species was performed using the Lipid Network Explorer with default settings (Köhler et al. 2021).

#### (2) RNASeq Pipeline and Gene Set Enrichment Analysis

A blocked experimental design with one replicate of each FFA in the library together with multiple BSA controls per 96-well plate was chosen (n=6). Raw data from NextSeq runs were de-multiplexed and converted to sample specific fastq files. Alignment was performed with STAR (Dobin et al. 2013), reads were counted with HTSeq (Anders, Pyl, and Huber 2015) and QC metrics were generated with RNA-SeQC (DeLuca et al. 2012).

The resulting count matrix was filtered by column for samples with more than 10^3^ detected genes (counts > 0) and by row for coding genes (as defined by the MGI database) with a row sum across all samples > 500 counts (with a total number of 500 samples). The resulting normalized and filtered count matrix was then variance stabilized using the vst method from the DESeq2 R package (Love, Huber, and Anders 2014). Surrogate variable analysis (SVA, R package)(Leek et al. 2012) was performed on the vst count matrix to account for linear batch effects. In addition, we performed differential expression analysis with DESeq2 for each sample, including derived surrogate variables to the linear model. To cluster the samples, we chose the top 500 most common significant differentially expressed genes (padj < 0.05) across the entire dataset. Samples were either transformed to z-scores or replicates were collapsed by calculating their signal to noise ratio (with respect to the BSA control) before performing hierarchical clustering based on Euclidean distance and Ward’s linkage method.

Clusters were extracted with the Dynamic Tree Cut function (Langfelder, Zhang, and Horvath 2008). After assigning each FFA to a cluster, we performed differential expression analysis based on cluster labels and BSA controls (based on the vst count matrix) and calculated adjusted p-values for each gene (Mann–Whitney U, Bonferroni). For gene set enrichment analysis (GSEA), gene lists for each FFA transcriptome were ranked by log2 fold changes as compared to BSA control to weigh genes according to their differential expression vs control. MsigDB H: HALLMARK gene sets, C2: KEGG and REACTOME gene sets and C5: GO BP gene sets were tested for enrichment (Liberzon et al. 2011; Subramanian et al. 2005). 250 Genesets with the most common, significant differential enrichment across the whole FFA library were selected and transformed into a normalized enrichment score (NES) matrix. Hierarchical clustering of those genesets resulted in the presented gene modules presented in Fig. 2A. Pearson correlation analysis of human islet gene expression (Taneera et al. 2012) was performed using MATLAB 2020a (MathWorks), as previously described (Brown et al. 2021). Genes significantly correlated with *CMIP* (FDR < 0.01,Benjamini-Hochberg) were subjected to KEGG pathway analysis using Metascape and visualized using Cytoscape.

#### (3) Rank-rank hypergeometric overlap (RRHO)

The R package RRHO2 (Cahill et al. 2018) was used to perform RRHO analysis to evaluate the cutoff-free overlap in differential expression results from FFA Clusters (C1- C5) versus human islets (Lytrivi, Ghaddar, et al. 2020) and mice pancreatic beta cells (Dusaulcy et al. 2019). RRHO2 plots show a heatmap with four quadrants to display the overlap between expression list comparisons. This includes: downregulated observations in both data sets (top right, Fig, S2E,F), downregulated in the comparative data and upregulated observations in our data (bottom right, Fig. S2E,F), upregulated in the comparative data and downregulated observations in our data (top left, Fig. S2E,F) and upregulated observations in both data sets (bottom left, Fig. S2E,F). For each comparison, one-sided enrichment tests were used on -log(p-values) with default step size for each quadrant.

#### (4) Structural analysis of FFAs

Molecular structural features were generated with the MOE software. A complete list of generated features are summarized in Table S5. Detailed descriptions of these features are available in the MOE user manual (v2016.08). The molecular feature matrix was filtered for non-constant features across the FFA library and transformed into z-scores. To decrease the linear dependence between features, meta-features were extracted from the original molecular features by hierarchical clustering based on Pearson correlation and Ward’s linkage method (cutreeDynamic, WGCNA, R package)(Langfelder, Zhang, and Horvath 2008). Clustering was performed iteratively until the maximum Pearson r correlation coefficient between any two meta-features was less than 0.8 (n=3 iterations). The first principal component of molecular feature clusters were used to define meta-features.

The random forest classifier (RFC) was performed (randomForest; R package)(Breiman 2001). Optimal values for ntree (number of trees to grow) and mtry (number of variables randomly sampled as candidates at each split) were determined empirically based on the classification accuracy of the RFC run on the entire dataset. The RFC was then run with leave-one-out cross validation. Meta-feature importance measures for RFC prediction were calculated (importance).

#### (5) Morphological feature analysis

Confocal images were acquired as described for Cell Painting using the Opera Phenix High Content Screening System (#HH14000000, Perkin Elmer)(Bray et al. 2016). Image analysis was performed using Harmony software (PerkinElmer). Single nuclei were first identified using Hoechst staining. Associated cell bodies around each nuclei were identified by the “find cytoplasm function” method A (individual cutoff 0.1) from the Harmony software in the 488 nm channel. ER and mitochondrial regions were identified (Find Image Region) based on their respective channels (stained with Concanavalin A and MitoTracker® respectively) For each cell, the standard morphological features (area and roundness); as well as advanced STAR (Symmetry, Threshold compactness, Axial or Radial) and SER (Spots, Edges, Ridges) morphology for ER and mitochondria were calculated. A full list of extracted features is summarized in Table S6. Data was exported per cell object, and downstream analysis was performed in R.

#### (6) MAGMA analysis pipeline, gene functional readout correlations

The MAGMA software (de Leeuw et al. 2015) (v 1.07) was used to perform SNP annotation, gene analysis to generate ranked lists of genes from GWAS summary statistics and gene set analysis (GSA) according to the instructions provided in the user manual. For multiple hypothesis testing we used a permutation-based approach to generate an empirical Null Hypothesis to account for the enrichment of beta cell genes in previous T2D GWAS analyses. For each gene set, we generated 1000 randomly sampled gene sets based on the MIN6 transcriptome of the same size and calculated the FDR accordingly.

### iPSC-beta Cell Differentiation

The iPSC-derived beta cells were differentiated as described previously (Maxwell and Millman 2021; Hogrebe et al. 2020) from an episomal reprogrammed iPSC line (Gibco,

#A18945). The factors added each day of differentiation can be found in Table S7. After the conclusion of the 28 day differentiation process, these cells were maintained in enriched serum-free media (ESFM) with media changes every other day. On day 30, the cultures were dissociated with TrypLE incubation for a maximum of 5 min, followed by neutralization with ESFM, TrypLE removal through centrifugation at 200xg for 3 min, filtration through a 40 µm filter to remove large clumps, and seeding onto HTB-9 ECM coated 96-well plates (Perkin Elmer, CellCarrier Ultra, #6055308) at 10,000/well.

### iPSC-microglia Cell Differentiation

AICS-0036-006 from the NIGMS Human Genetic Cell Repository at the Coriell Institute for Medical Research was used in this study. iPSCs were cultured in Essential 8 (E8) (Thermo Fisher Scientific) media on Matrigel (Corning) coated 6-well plates. Media was changed daily until confluence. iMGLs were differentiated as previously described (Abud et al. 2017; Dolan et al. 2022). When confluent, iPSCs were dissociated using Accutase (Stem Cell technologies), centrifuged for 5 mins at 300xg and counted using trypan blue (Thermo Fisher Scientific). 200,000 cells/well were resuspended in E8 containing 10 μM Y27632 ROCK inhibitor (Selleckchem) in low adherence 6-well plates (Corning). For the first 10 days, cells were cultured in HPC medium [50% IMDM (Thermo Fisher Scientific), 50% F12 (Thermo Fisher Scientific), ITSG-X 2% v/v (Thermo Fisher Scientific), L-ascorbic acid 2-Phosphate (64 ug/ml, Sigma), monothioglycerol (400mM, Sigma), Poly(vinyl) alcohol (PVA) (10mg/ml, Sigma), Glutamax (1X, Thermo Fisher Scientific), chemically-defined lipid concentrate (1X, Thermo Fisher Scientific) and non-essential amino acids (Thermo Fisher Scientific)]. At day 0, embryoid bodies (EB) were gently collected, centrifuged at 100xg and resuspended in HPC medium supplemented with 1 μM ROCK inhibitor, FGF2 (50 ng/ml, Thermo Fisher Scientific), BMP4 (50ng/ml, Thermo Fisher Scientific), Activin-A (12.5ng/ml, Thermo Fisher Scientific) and LiCL (2 mM, Sigma), then incubated in a hypoxic incubator (5% O2, 5% CO2, 37 °C). On day 2, cells were gently collected and the media changed to HPC medium supplemented with FGF2 (50ng/ml, Thermo Fisher Scientific) and VEGF (50 ng/ml, PeproTech) and returned to the hypoxic incubator. On day 4, cells were collected and media changed to HPC medium supplemented with FGF2 (50ng/ml, Thermo Fisher Scientific), VEGF (50 ng/ml, PeproTech), TPO (50 ng/ml, PeproTech), SCF (10ng/ml, Thermo Fisher Scientific), IL6 (50ng/ml, PeproTech) and IL3 (10ng/ml, PeproTech) and incubated in a normoxic incubator (20% O2, 5% CO2, 37C). At day 6 and 8, 1 ml of day 4 media was added in each well. On day 10, cells were collected, counted using trypan blue and frozen in Cryostor (Sigma Aldrich) in aliquots of 300,000-500,000 cells.

For iMGL differentiation, cells were thawed, washed 1x with PBS and plated at 200,000 cells per well in 6-well plates coated with matrigel in iMGL media [DMEM/F12 (Thermo Fisher Scientific), ITS-G (2% v/v, Thermo Fisher Scientific), B27 (2% v/v, Thermo Fisher Scientific), N2 (0.5% v/v, Thermo Fisher Scientific), monothioglycerol (200 mM, Sigma), Glutamax (1X, Thermo Fisher Scientific), non-essential amino acids (1X, Thermo Fisher Scientific)] supplemented with M-CSF (25 ng/ml, PeproTech), IL-34 (10 ng/ml, PeproTech) and TGFB-1 (50ng/ml, PeproTech). Cells were fed every 2 days and replated at day 22. On day 30, cells were collected and replated in iMGL media supplemented with M-CSF (25 ng/ml, PeproTech), IL-34 (10 ng/ml, PeproTech), TGFB- 1 (50ng/ml, PeproTech), CD200 (100ng/ml, VWR) and CX3CL1 (100ng/ml, PeproTech).

### CMIP KO Clone Selection

Lentivirus containing a plasmid programmed to express either CMIP-specific sgRNA (ACGTCTTCAATGGCGCTGTAGG, Millipore Sigma, Sanger Clone MM5000005403) or non-targeting control sgRNA (Millipore Sigma, CRISPR20-1EA) was obtained through transfecting HEK293T cells with these plasmids in combination with a second generation CMV lentiviral packaging system and FuGENE transfection agent (E2311, Promega).

MIN6 cells stably expressing Cas9 were infected with this lentivirus at half volume with 8 μg/ ml polybrene (TR-1003-G, Sigma) overnight. The media was changed 12 h later, and 48 h after the first media change, 1 µg/ml of puromycin was added to the media. After one week of puromycin selection, the cultures were dispersed into single cells, seeded into 96-well plates for expansion, and sequenced for CMIP mutations. One WT clone and one possessing a truncation within the first 15% of the longest CMIP isoform were used for the experiments described.

To derive CMIP KO cells from the human iPSC-beta cell line, we delivered Cas9- RNP complexes with either CMIP-specific sgRNA or non-targeting sgRNA (same as Min6 above) to the parental iPSC cells using a Lonza nucleofector kit. Upon confirmation of successful transfection and generation of a stable knockout line, single clones were isolated and sequenced to ultimately identify the clone used in the studies described.

### CMIP Overexpression for Rescue Studies

To re-express CMIP back into the MIN6 CMIP KO line, we obtained lentivirus from VectorBuilder containing a mouse CMIP ORF (NM_001163262.1) with a modified PAM site under a CMV promoter and accompanied by a neomycin resistance gene. We used the same vector with amino acids 2-83 of E. coli beta-galactosidase as a substitute for the CMIP ORF for our control line. 600,000 cells were seeded into each well of a 6-well plate and one week later were infected with up to 100ul virus per well (and 8 ug/ml polybrene (TR-1003-G, Sigma)) overnight. The media was changed 12 h later, and 48 h after the first media change, 800 µg/ml of G418 was added to the media. After selection for one week, the cells were used for the experiments described.

### Cell Viability

For the high throughput cell viability assay, cells were seeded in 384-well plates (Perkin Elmer, CellCarrier Ultra, #6057300) and treated for 24, 48 and 72 h with the FFA library (n=7 / FFA). Just before readout, cell nuclei were stained with Hoechst (Thermo Fisher Scientific) for 1 h at 37 °C and imaged with the Opera Phenix High Content Screening System (#HH14000000, Perkin Elmer). Number of counted nuclei was determined with the image analysis software Harmony (PerkinElmer) and used as a proxy for cell viability. For validation experiments, cells were treated for 48 h with representative FFAs in CellCarrier-384 Ultra Microplates. Caspase 3/7 (Thermo Fisher Scientific, #C10423) activation and propidium iodide (Thermo Fisher Scientific, #P3566) staining were used to calculate the fraction of apoptotic cells and dead cells, respectively. Single cells were identified and counted after staining their nuclei with Hoechst. Fluorescence intensities were measured and the threshold for caspase 3/7 and propidium iodide positive staining was determined manually. Cell viability was calculated as the fraction of cells that were neither caspase 3/7 nor propidium iodide positive. For the EA dose-response curve, MIN6 cells were grown as described and then treated for 65 hours with BSA, EA or OA. The concentrations were (in mM): 0.7, 2.1, 6.2, 18.5, 55.6, 166.7, 500, 1000 and were prepared by serial dilution from a stock concentration. Cell viability was assessed as described.

#### iPSC-derived microglia

On day 30 of the differentiation, iMGLs were plated in 96-well plates (Perkin Elmer, CellCarrier) at a density of 20,000 cells per well. One day before treatment, FFA and BSA were reconstituted in iMGL media and gently rocked overnight at room temperature. At day 40, iMGL were treated with a final concentration 250 μM of each FFA or BSA for 24 h followed by live imaging. Cells were imaged using the Opera Phenix confocal system using a 20X objective, temperature was maintained at 37 °C and CO2 at 5% during the imaging period. One plane was taken per picture and 20 images were taken per well. Harmony software was used for analysis, flatfield correction (Basic Method) and cell identification using EGFP signal (common Threshold 0.02; area >50 µm^2^). Total number of cells per well was used for quantification with an n=4 for all conditions.

#### Kidney tubular epithelial cells

Prior to treatment, BSA-bound FFAs were reconstituted and incubated overnight in RenaLife media at room temperature. Cells were seeded in CellCarrier-384 Ultra Microplates and maintained at 37°C with 5% CO2. Cells were treated with 500 µM FFA or BSA for 15 h followed by live imaging using the Opera Phenix High Content Screening System. Number of cells and number of dead cells was determined on Harmony image analysis software using digital phase contrast and propidium iodide staining (Thermo Fisher Scientific, #P3566), respectively. Viability was assessed by calculating the number of propidium iodide-negative cells per total number of cells per well (n=4).

### Western Blot

MIN6 cells were lysed (#9803, Cell Signaling Technology) in the presence of protease inhibitors (#05892791001, Roche) and phosphatase inhibitors (#04906837001, Roche). Protein concentrations were quantified with the Pierce BCA Protein Assay Kit (#23225, Thermo Fisher Scientific). NuPAGE LDS sample buffer (#NP0008, Thermo Fisher Scientific) was added to normalized protein lysates together with NuPAGE reducing agent (#NP0004, Thermo Scientific). Lysates were heated to 95 °C for 5 min prior to SDS-PAGE gel electrophoresis (NuPAGE MES SDS running buffer, Thermo Fisher Scientific, #NP0002). Proteins were transferred to a nitrocellulose membrane (#1704158, BioRad) with the Trans-Blot® Turbo TM Blotting System (#1704155, BioRad) according to the manufacturer’s protocol. Membranes were blocked in 5% Nonfat Dry Milk (#9999S, Cell Signaling Technology) in PBS with 0.1% Tween® 20 (PBS-T). AKT and pAKT blots were blocked instead in 5% Bovine Serum Albumin (#1900-0016, LGC Clinical Diagnostics) in PBS-T. Primary antibodies were incubated at 4 °C overnight, secondary antibodies were incubated at room temperature for 1 h. Super Signal West Dura (#34076, Thermo Fisher Scientific) or SuperSignal West Pico (#34087, Thermo Fisher Scientific) were used to visualize immunoreactive bands imaged by G:BOX Chemi XT4 (BOX-CHEMI-XT4, Syngene). Primary antibodies used in this study were CPT1A: (#ab128568, Abcam), ATF4: (#11815, Cell Signaling Technology), CHOP: (#2895, Cell Signaling Technology), CMIP: (NBP2-58180, Novus Bio), AKT: (#9272, Cell Signaling Technology), pAKT: (#4060, Cell Signaling Technology), AMPKL (#2532, Cell Signaling Technology), pAMPKL (#2535, Cell Signaling Technology), FOXO1: (#2880, Cell Signaling Technology), pFOXO1: (#9464, Cell Signaling Technology), GAPDH-HRP: (#3683, Cell Signaling Technology). Secondary antibodies used were: anti-rabbit IgG-HRP (#7074, Cell Signaling Technology), anti-mouse IgG-HRP (#7076, Cell Signaling Technology).

### Coimmunoprecipitation (co-IP)

Two 10cm dishes of Min6 cells were lysed with 1ml of co-IP lysis buffer (100mM NaCl, 5mM EDTA, 50mM Tris-HCl pH 7.5, 1% NP-40, protease inhibitor tablet, phosphatase inhibitor tablet) after being washed once with ice cold PBS. The lysate was rotated at 4°C for 30m, spun at 13,000xg for 20m at 4°C, and the supernatant was collected. 1mg of protein was combined with 10ug of either CMIP antibody (12851-1-AP, Proteintech) or rabbit IgG control antibody (10500C, Thermo Fisher) and rotated overnight at 4°C. The following day, Pierce Protein A/G Magnetic Beads (88802, Thermo Fisher) were warmed to room temperature, washed twice with 1ml TBS-T (150mM NaCl, 50mM Tris-HCl pH 7.5, 0.05% Tween-20) on a magnet, and resuspended in the original volume of co-IP lysis buffer. 25ul of beads were added to each lysate tube and the solution was rotated at 4°C for 1h. The beads were then washed 3x with 1ml TBS-T, once with 1ml distilled water, and eluted with 30ul of 2X NuPAGE LDS Sample Buffer and 1X NuPAGE Sample Reducing Agent (DTT). The beads were incubated at room temperature for 10m with occasional flicking of the tube, and then the supernatant was extracted on the magnet. The supernatant was incubated at 95°C for 10m and then the samples were run on a gel as detailed in the Western Blot section above. The primary antibodies used for staining were CMIP: (12851-1-AP, Proteintech) and PI3K p85L (#4292, Cell Signaling Technology).

### Immunofluorescence (IF) staining

#### MIN6 NFkB Immunofluorescence

MIN6 cells grown on 384-well CellCarrier Ultra microplates (#6057308, PerkinElmer) were fixed for 10 min in PBS containing 4% PFA (Electron Microscopy Sciences), permeabilized for 15 min in 0.5% Triton X-100 (Sigma-Aldrich), blocked for 1 h in blocking reagent (100 mM Tris HCL pH 8; 150 mM NaCL; 5 g/L Blocking Reagent (#11096176001, Roche)) and treated for 1.5 h with primary antibody diluted in blocking reagent (NF-κB p65/RELA, Rabbit monoclonal antibody, 1:200, #8242, Cell Signaling Technology). Cells were washed three times in PBS and incubated for 0.5 h with fluorescent-labeled secondary antibody in blocking solution (1:500, Alexa Fluor 568 Goat anti-Rabbit IgG, (#A11036, Thermo Fisher Scientific)). Cytoplasmic actin filaments were stained with Phalloidin conjugated with Alexa 647 (1:40, #A22287, Thermo Fisher Scientific) and nuclei were counterstained with Hoechst (1:2000, #H3570, Thermo Fisher Scientific). Cells were washed three times in PBS and imaged using the Opera Phenix High Content Screening System (#HH14000000, Perkin Elmer). A minimum of nine fields were acquired per well using 20x water immersion objectives in confocal mode. Image analysis was performed using the Harmony software (PerkinElmer). Cell nuclei were first identified using Hoechst staining and a nuclear region was defined for each cell. Phalloidin staining was then used to detect and define the cytoplasmic region of the cell. RELA fluorescent intensity was measured separately in the nuclear and cytoplasmic regions and a threshold for a nuclear translocation was defined using negative (BSA) and positive (TNF) controls. For each well, the fraction of cells identified for RELA nuclear translocation was calculated.

#### MIN6 LC3B Immunofluorescence

MIN6 cells grown on 384-well CellCarrier Ultra microplates (#6057308, PerkinElmer) were treated for 48 h with 500 µM FFAs or 25 nM of rapamycin (Sigma-Aldrich, R8781) or bafilomycin A1 (Sigma-Aldrich SML1661). Cells were fixed for 20 min in ice-cold methanol (Sigma-Aldrich, 154903), washed twice with PBS, permeabilized for 15 min in 0.5% Triton X-100 (Sigma-Aldrich, 10789704001), washed twice with PBS, blocked for 1 h in 5% BSA in PBS (SeraCare, 1900-0016) and incubated for 1.5 h with primary antibody diluted in blocking reagent (Lc3b (D11) XP® Rabbit mAb, 1:500, #3868, Cell Signaling Technology). Cells were washed four times in PBS and incubated for 45 min with fluorescent-labeled secondary antibody in blocking solution (1:500, Alexa Fluor 568 Donkey anti-Rabbit IgG, (#A10042, Thermo Fisher Scientific)). Nuclei were counterstained with Hoechst (1:2000, #H3570, Thermo Fisher Scientific). Cells were washed four times in PBS and imaged using the Opera Phenix High Content Screening System (#HH14000000, Perkin Elmer). A minimum of nine fields were acquired per well using 63x water immersion objectives in confocal mode. Image analysis was performed using the Harmony software (PerkinElmer). Cell nuclei were first identified using Hoechst staining and a nuclear region was defined for each cell. LC3B puncta were then detected based on intensity and size with rapamycin and bafilomycin treatments serving as controls.

#### Primary Human Islet Immunofluorescence

Primary human islets from three independent cadaveric donors were received from the Integrated Islet Distribution Program. Intact islets were dissociated using accutase followed by trituration and stained using Trypan Blue to confirm viability. They were seeded at 15,000/well into 384-well Cell Carrier Ultra microplates (Perkin Elmer, CellCarrier Ultra, #6057300) coated with HTB-9-derived ECM, and treated two days later with FFAs at 250 µM, 500 µM, and 1 mM. After treatment for 5 days, the islets were fixed for 20 min in 3% PFA followed by permeabilization for 20 min with 0.2% TritonX-100. Blocking with 2% BSA in PBS (SeraCare, AP-45100-80) was conducted for 1 h at room temperature, followed by primary incubation with c-peptide antibody (Developmental Studies Hybridoma Bank at the University of Iowa, GN-ID4) at 1:100 overnight at 4 °C. The plate was washed 3x with PBS followed by secondary incubation with 568 goat anti-rat (Life technologies, A11077) at 1:1000 and Hoechst (1:2000, #H3570, Thermo Fisher Scientific) for 1 h at room temperature. The plate was then washed 5x with PBS and imaged on the Opera Phenix High Content Screening System (#HH14000000, Perkin Elmer). A minimum of nine fields were acquired per well using a 20x water immersion objective in confocal mode. Image analysis was performed using the Harmony software (PerkinElmer). Beta cells were identified and counted by c-peptide positive staining. All three donors displayed similar trends in c2 toxicity; the data presented in Fig. 5C are representative of all donors.

#### iPSC-Derived Beta Cell Immunofluorescence

IPSC-derived beta cells after 28 days of differentiation as per the protocol listed above were dissociated and seeded at 10,000/well into 96-well Cell Carrier Ultra microplates (Perkin Elmer, CellCarrier Ultra, #6055308) coated with HTB-9-derived ECM. These cells were treated three days later with FFAs at 250 µM, 500 µM, and 750 µM for 24, 48, and 72 h followed by fixation with 4% PFA for 30 min at room temperature. The plates were washed twice with PBS, permeabilized for 15 min with 0.5% Triton-X 100, washed twice again with PBS, and blocked with 5% BSA in PBS for 1 h at room temperature. Hoechst was added (1:2000, #H3570, Thermo Fisher Scientific) and incubated for 1 h at room temperature. The plate was then washed thrice with PBS and imaged on the Opera Phenix High Content Screening System (#HH14000000, Perkin Elmer). A minimum of nine fields were acquired per well using a 20x water immersion objective in confocal mode. Image analysis was performed using the Harmony software (PerkinElmer) and the cells were counted.

### ER Calcium Levels

MIN6 cells were plated in 384-well plates (Aurora, Black 384 SQ Well 188 micron Film, #1022-10110) and treated with the FFA library for 24 h prior to readout (n=5 / FFA). Cells were carefully washed three times with HBSS (with calcium, Thermo Fisher Scientific, #14025076) using an automated simultaneous pipettor (analytikjena CyBio^®^ Well vario) and incubated with the fluorescent calcium indicator Fluo4 (2 µM, Life Technologies, #F14202) in DMEM without supplementation for 1 h at room temperature. Then, cells were washed again in HBSS (with calcium) and incubated for another 30 min at room temperature in DMEM without supplementation. Just before the readout, cells were washed in calcium free assay buffer solution (140 mM NaCl, 5 mM KCl, 10 mM HEPES, 2 mM MgCl2, 10 mM EGTA, 10 mM Glucose) and left with 25 µl assay volume per well. Assay plates were immediately transferred to the FLIPR Tetra® High- Throughput Cellular Screening System. The plate was recorded with a frequency of 1Hz for 10 min. Baseline was recorded for 30 s before the automated liquid transfer system of the FLIPR added the SERCA inhibitor Thapsigargin (final concentration 10 µM) in calcium free assay buffer. The resulting passive efflux of calcium from the ER induced a transient cytosolic fluorescence signal and the peak amplitudes were used to indirectly quantify ER calcium levels (See Fig. S3A). The resulting trajectories were corrected for a pipetting artifact and baseline normalized. Log2 Fold changes were calculated according to plate location specific negative (BSA) controls. We allowed for the exclusion of one outlier / FFA / plate (n=5) based on a 3-sigma cutoff. P-values were calculated with Student’s *t*-test (two-sided) and corrected for multiple testing (Benjamini & Hochberg).

### Glucose Stimulated Insulin Secretion (GSIS)

To measure GSIS, cells were plated in 96-well plates (Perkin Elmer, CellCarrier Ultra, #6055308) and treated with the FFA library for 24 h prior to readout (n=6 / FFA). First, cells were pre-incubated in Krebs Ringer Buffer (KRB) with 2.8 mM glucose for 1 h and then stimulated with fresh KRB containing 16.7 mM glucose for another hour. The supernatant was then transferred to a mouse insulin ELISA (Thermo Fisher, EMINS) at a dilution of 1:150. The cells were imaged on the Opera Phenix High Content Screening System, (#HH14000000, Perkin Elmer) and subsequently counted via Hoechst nuclear staining using Harmony software (PerkinElmer). A minimum of four fields were acquired per well using a 10x air objective in a confocal mode. The insulin concentrations measured via ELISA were normalized to the number of cells in each respective well.

### Membrane Fluidity Assay

This assay has been well established and optimized in INS-1E cells, a rat beta cell line. INS-1E cells were seeded at 20,000 cells per well in a CellCarrier Ultra 96-well plate (Perkin Elmer). Cells were incubated for 24 h, washed once with PBS, treated with BSA-bound fatty acids in a serum-free media and further incubated for 18 h at 37°C. Cells were washed twice with PBS and stained with Laurdan dye (6-dodecanoyl-2- dimethylaminoaphthalene) (Thermofisher) at 10 µM for 45 min. Cells were washed twice with PBS and images were acquired using an Opera Phenix High-Content Screening system (Perkin Elmer). Temperature (37°C) and carbon dioxide (5%) were controlled during live-cell imaging. Cells were excited with a 405 nm laser and the emission recorded between 435 and 480 nm (ordered phase) and between 500 and 550 nm (disordered phase). Images were analyzed using Harmony High-Content Imaging and Analysis Software (Perkin Elmer) and data represented as general polarization (GP) index as calculated by GP = (intensityorder - intensitydisorder)/(intensityorder + intensitydisorder) (Harris, Best, and Bell 2002; Parasassi et al. 1991).

## Supplemental Figure Legends

**Fig. S1.**
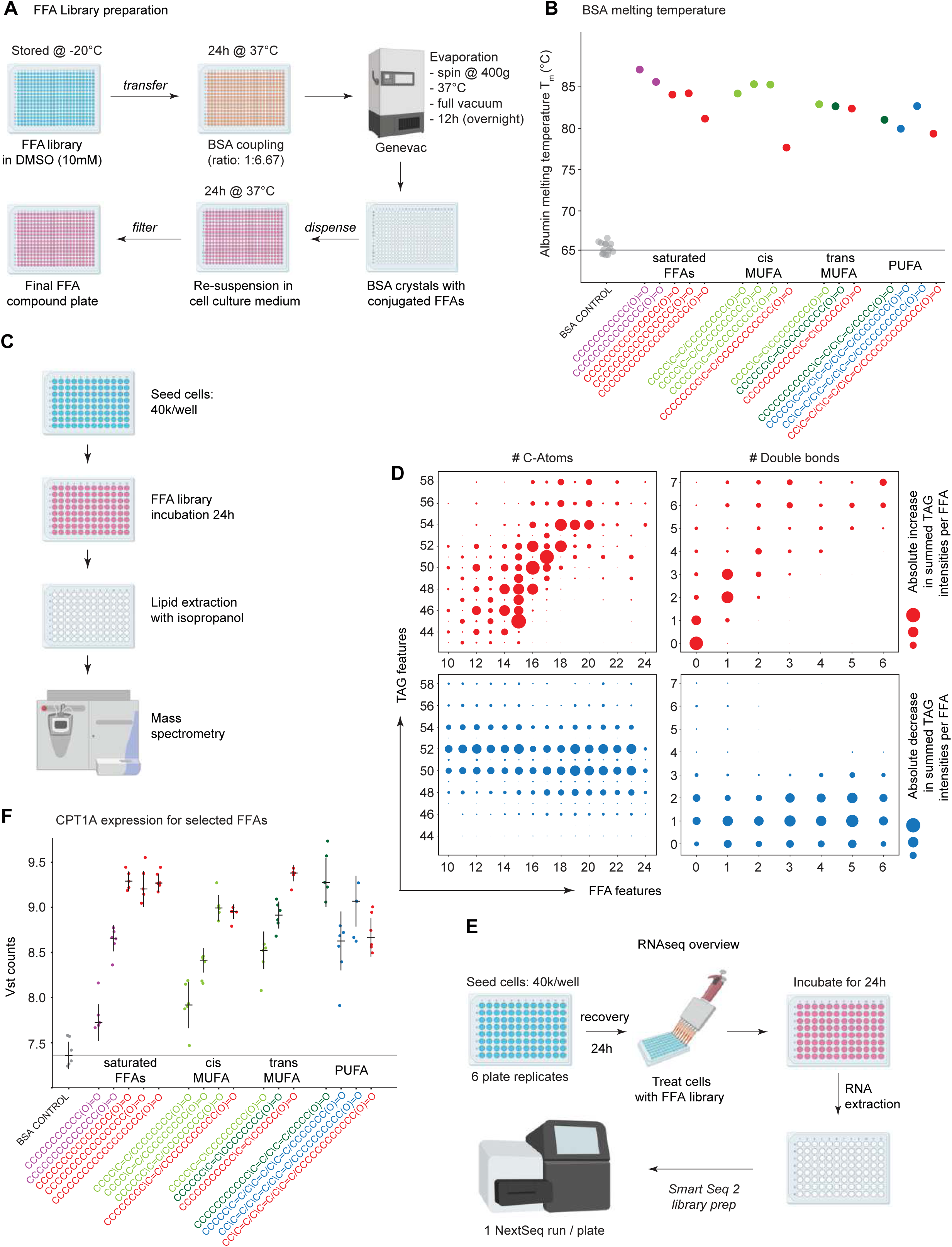
(A) Protocol overview describing major steps of the procedure for preparing FFAs. (B) Shift in bovine serum albumin (BSA) melting temperature Tm (y-axis). After library preparation, structurally representative FFAs (structure SMILES, x-axis) consistently increased Tm confirming successful BSA:FFA loading. Colors represent cluster identities (Fig. 1D). (C) Overview of sample preparation for lipidomics experiment. (D) Lipidomic analysis indicated that cells exposed to the library of 61 externally applied FFAs incorporated them into their TAG fraction as detected by mass spectrometry. Qualitative correlation of structural features (number of C atoms, number of double bonds) of externally applied FFAs (x-axis) versus structural features of triglycerides (TAGs, y-axis) detected by lipidomics (methods). Upper two panels (red dots) show absolute increases in summed triglyceride intensities per FFA, lower two panels (blue dots) show absolute decreases in summed triglyceride intensities per FFA (negative control). (E) RNAseq protocol overview. (F) CPT1A expression for selected FFAs. Consistently significant differential expression (padj < 0.05) confirmed successful delivery of FFAs to the cells.

**Fig. S2.**
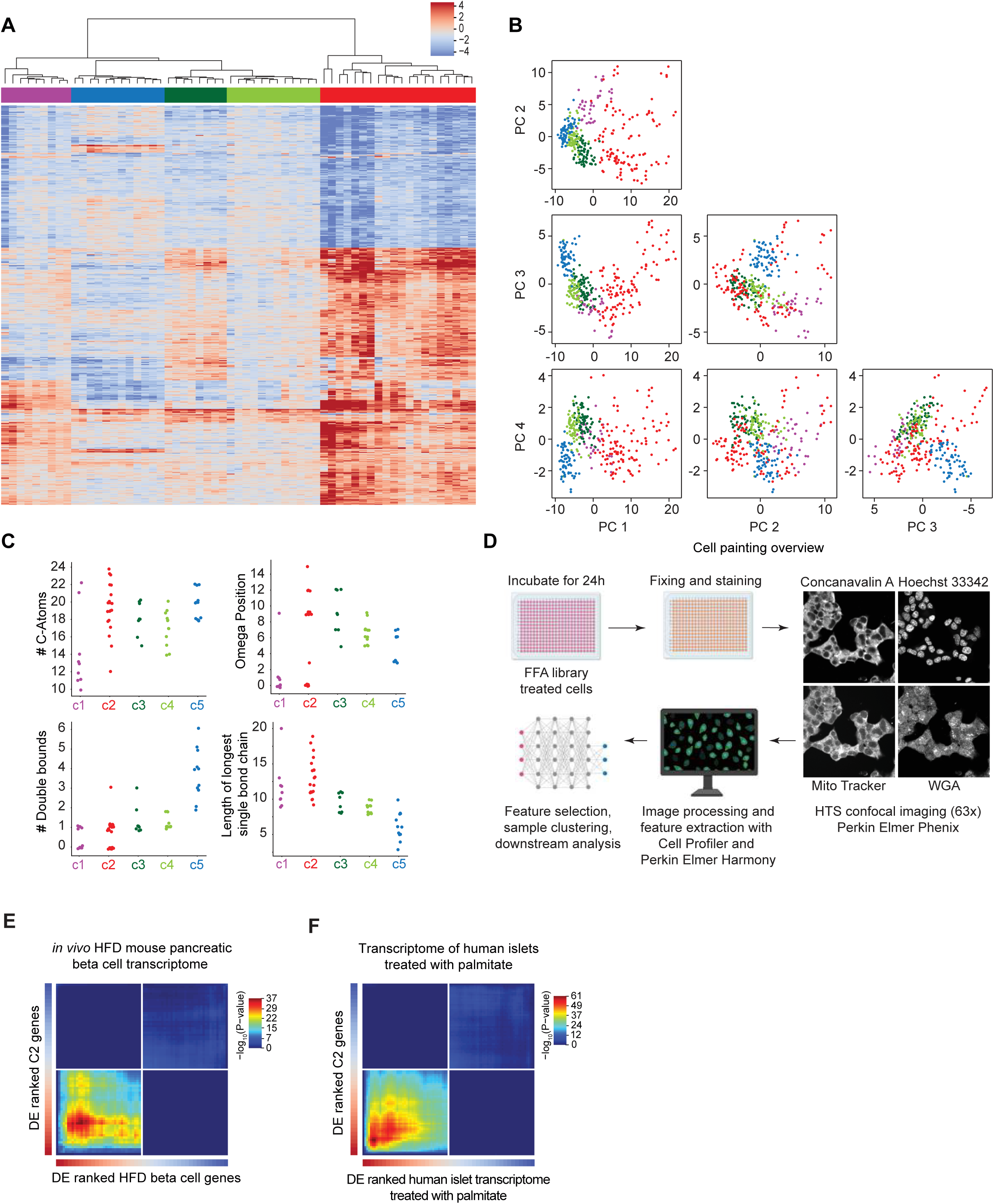
(A) Hierarchically clustered heatmap based on z-scores of top 500 differentially expressed genes across the entire dataset (collapsed replicates). Clusters were extracted with the Dynamic Tree Cut function (Langfelder, Zhang, and Horvath 2008). (B) Corner plot of first four Principal Components (PC) of all replicates, capturing 75% of the total variance in the dataset. Colors represent FFA cluster identities (as in Fig. 1D). (C) The structural characteristics of each FFA including number of carbon atoms, number of double bonds, omega position (distance from omega carbon to first double bond), and length of the longest single bond chain are mapped across each cluster. Cluster number and color corresponding to that seen in Fig. 1D are along the x-axis; structural characteristic distribution of FFAs within each cluster is found along the y-axis. (D) Overview of the Cell Painting method. (E,F) Rank-rank hypergeometric overlap (RRHO) maps show significant overlap of the C2-derived lipotoxic- ity signature with published transcriptomes of pancreatic beta cells isolated from mice fed a high fat diet (E) and human islets exposed to palmitate (F).

**Fig S3.**
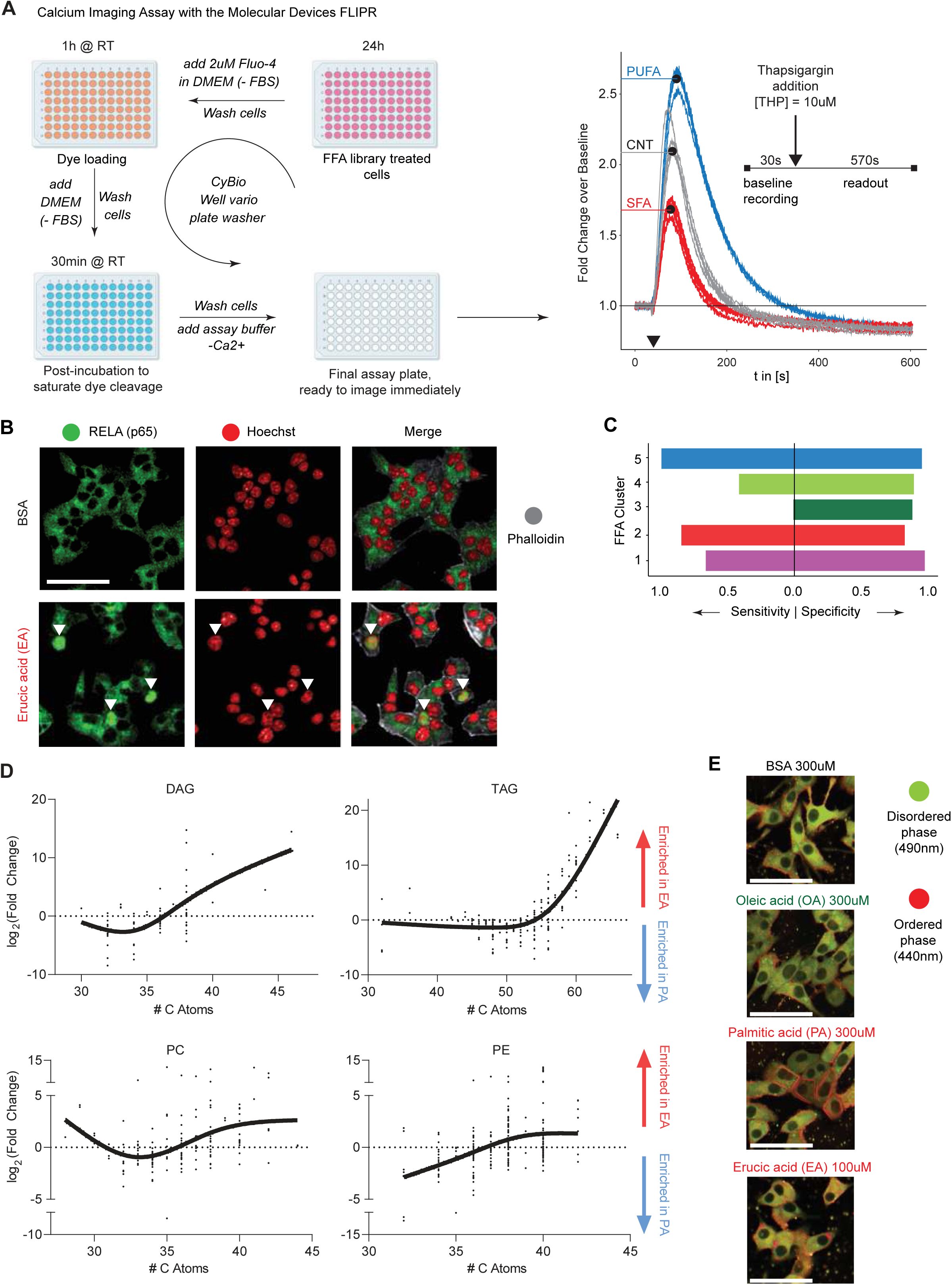
(A) Overview of the Ca2+ imaging protocol used to quantify dynamic changes in cytosolic Ca2+ levels. Right: Representative example trajectories (n = 5 replicates) of thapsigargin-stimulated (10 mM, black arrow) Ca2+ signals, as fold changes over baseline (y-axis). The inset outlines the measurement protocol. (B) Representative images from immunofluorescence of nuclear translocation of RELA (green) upon exposure to EA (t = 18 h). BSA, negative control. Nuclei were stained with Hoechst (red) and the cytoplasmic cytoskeleton with phalloidin (grey). Complete translocation of RELA into the nucleus (white arrows) was detected in EA-treated cells, highlighted in the merged image. Scale bars: 100 μm. (C) Decision tree prediction accuracy (sensitivity and specificity) of assigning FFAs into transcriptomically defined clusters based on a structural meta-feature matrix (see methods). (D) Log-fold changes in the abundance of specific lipid species between PA-treated and EA-treated samples. Species above the x-axis are enriched in EA-treated samples, and species below are enriched in PA-treated samples. DAG, diglycerides; TAG, triglycerides; PC, phosphatidylcholines; PE, phosphatidylethanolamines. (E) Representative images of INS1E beta cells assayed in Figure 5G highlighting the disordered phase (green) and the ordered phase (red) of the Laurdan dye. Scale bars: 50 μm.

**Fig. S4.**
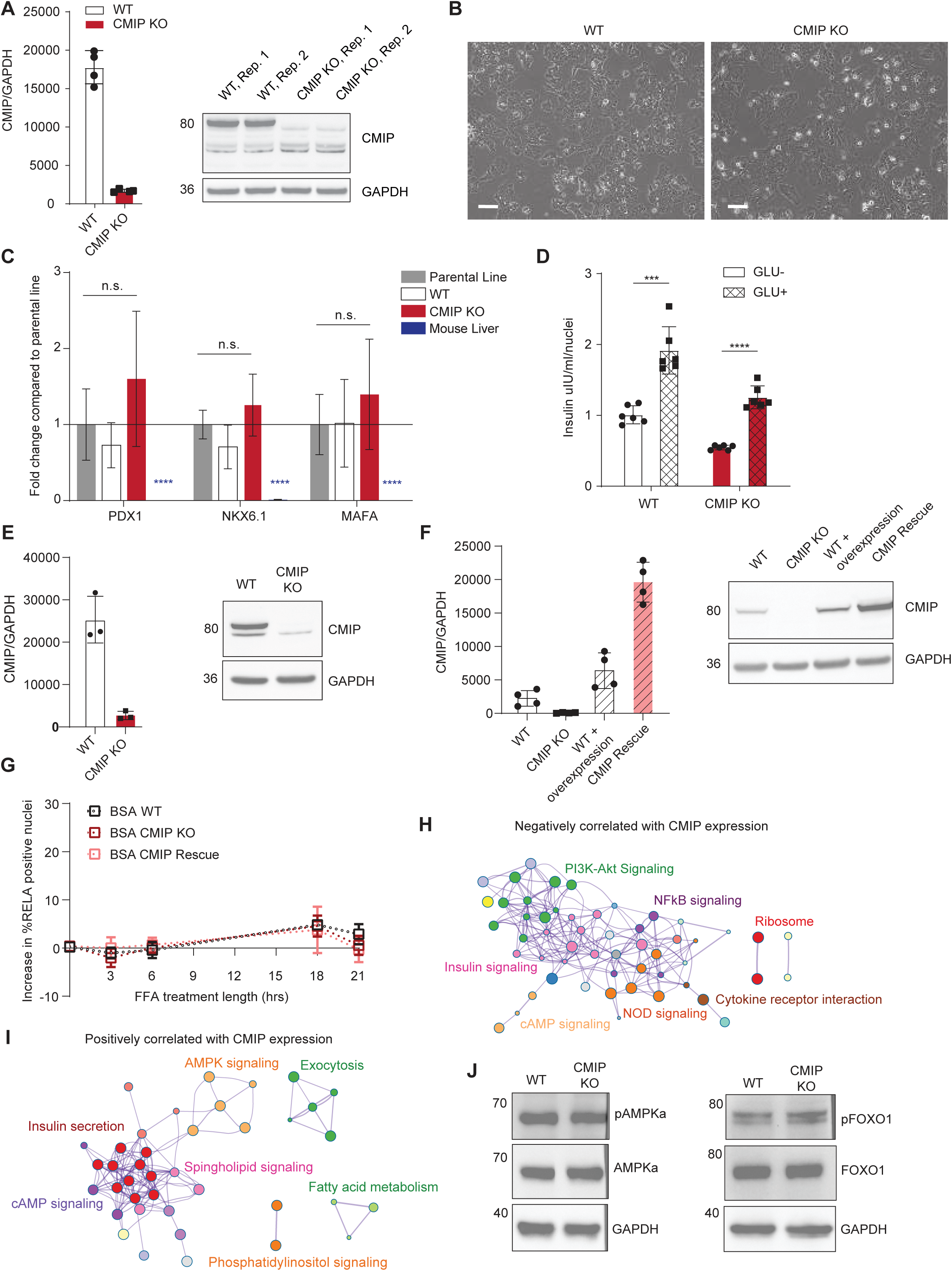
(A) Quantification of CMIP protein expression in CMIP KO cells normalized to GAPDH (n = 4 blots). The primary endogenous CMIP isoform was fully deleted in the CMIP KO cells. (Right) Western blot of WT and CMIP KO lysates for CMIP and GAPDH (loading control). (B) 10x Bright- field images of WT and CMIP KO cells show no morphologic differences. Scale bars: 100 μm. (C) WT and CMIP KO cells expressed similar mRNA levels of beta cell markers compared to the parental line, and expressed significantly higher levels of PDX1, NKX6.1, and MAFA compared to the mouse liver cDNA (negative control). Two-sided ANOVA analysis, ****p < 0.0001, n = 3 replicates. (D) Insulin concentration in the supernatant of the WT and CMIP KO cells after treatment with low glucose KRB buffer (2.8 mM) or high glucose buffer (16.7 mM) for one hour each. The concentration was normalized to the number of cells present in each well. Insulin secretion doubled upon incubation with high glucose buffer for both WT and CMIP KO cells (n=6 wells). Data are mean ± SD. Multiple t-test (two-sided), ***p < 0.001, ****p < 0.0001, with Bonferroni correction. (E) Quantification of CMIP protein expression in each human iPSC-derived beta cell line normalized to GAPDH (n=3 blots). The primary endogenous CMIP isoform is fully deleted in the CMIP KO line. (Right) Western blot of WT and CMIP KO iPSC lysates for CMIP and GAPDH (loading control). (F) Rescue of CMIP expression in MIN6 CMIP KO cells by overexpression of the most abundant CMIP isoform. Introduction of the overexpression plasmid successfully induces expression of a CMIP isoform at the appropriate molecular weight as assessed by Western blot (right). GAPDH, loading control (n=3 blots). (G) Reintroduction of CMIP into CMIP KO cells does not alter baseline NFkB activation. Percentage of cells with RELA nuclear translocation (nuclear to cytoplasmic signal > 1.2) upon treatment with BSA (500 μM) at 0, 3, 6, 18, and 21 h after subtraction of baseline signal in non-treated cells. Data are mean ± SD, n = 6-9 replicates/timepoint. (H,I) Network analysis of human pancreatic islet transcriptome data (GSE38642; n=51 donors; (Taneera et al. 2012)) illustrating the interaction between enriched KEGG pathways from genes negatively (H) and positively (I) correlating with CMIP expression (FDR < 0.01). (J) CMIP KO does not alter AMPKɑ or FOXO1 activation. On the left is a series of Western blots for pAMPKɑ, AMPKɑ, and GAPDH (loading control) indicating no difference between the WT and CMIP KO cells (n=2 blots). On the right is a series of Western blots for pFOXO1, FOXO1, and GAPDH (loading control) indicating no difference between the WT and CMIP KO cells (n=2 blots).

